# APOE4 expression confers a mild, persistent reduction in neurovascular function in the visual cortex and hippocampus of awake mice

**DOI:** 10.1101/2021.05.26.445731

**Authors:** Orla Bonnar, Kira Shaw, Dori M. Grijseels, Devin Clarke, Laura Bell, Silvia Anderle, Sarah L. King, Catherine N. Hall

## Abstract

Young mice possessing the e4 allele of the Apolipoprotein (APOE) gene (a risk factor for Alzheimer’s disease (AD)) have previously been shown to have dramatic decreases in vascular function, suggesting APOE4 may confer its risk via the vasculature. However, in human carriers, vascular and cognitive function decrease later in life. Mouse data may be confounded by an increased impact of anaesthesia and surgery in APOE4 animals, and has also focused on sensory cortices, ignoring medial lobe structures more sensitive to AD.

To clarify how APOE4 expression alters neurovascular function, we studied the visual cortex and hippocampus of awake APOE3 and APOE4 targeted replacement mice, using 2-photon microscopy of neurons and blood vessels.

We found milder vascular deficits than studies using anaesthetised preparations: functional hyperaemia was unaffected in APOE4 mice and neuronal or vascular function did not decrease with age. Instead, vascular responsiveness was lower at all ages, arteriole vasomotion was reduced and neuronal calcium signals during visual stimulation were increased. This suggests that, independently, APOE4 expression is not catastrophic but alters neurovascular physiology towards a state more sensitive to insults such as surgery or beta amyloid accumulation. Understanding how APOE4 expression interacts with these insults will be critical for understanding the emergence of AD in APOE4 carriers.

## Introduction

The brain’s dense network of vasculature performs a myriad of roles that are crucial for neuronal health and function, including the fine-tuned delivery of oxygen and glucose via the blood, in response to neuronal activity (neurovascular coupling), and the clearance of waste products from the brain, including beta amyloid (Aβ). Both these processes require controlled constriction and dilation of microvascular smooth muscle cells and pericytes. In neurovascular coupling, this likely matches blood flow to energy demand (Iadecola, 2017), while perivascular clearance along blood vessels is promoted by slow (∼0.1Hz) oscillations of arterioles, termed vasomotion (Aldea et al., 2019; van Veluw et al., 2020). Alterations to these processes are associated with several neurodegenerative diseases, including Alzheimer’s disease (AD). In AD, these changes in cerebral blood flow likely establish a vicious cycle that exacerbates AD pathology, whereby decreased blood flow increases Aβ accumulation by promoting its formation in hypoxic conditions. Blood flow is then limited by accumulating Aβ, as blood vessels are constricted by soluble Aβ, and accumulation of Aβ within vessels likely inhibits vasomotion (Di Marco et al., 2015; Hamilton et al., 2010; Kisler et al., 2017; Sun et al., 2006; van Veluw et al., 2020). Vascular dysregulation has been shown to be among the earliest pathologies found in AD (Iturria-Medina et al., 2016), so disrupted blood flow may be the primary cause of emerging AD pathology.

Recent work from our group has demonstrated heterogeneity in vascular function across brain regions, where CA1 region of the hippocampus was less able to match blood supply to neuronal demand compared to the primary visual cortex (Shaw et al., 2021a). Such regional variability follows not only the known differences in vulnerability to hypoxia (Michaelis, 2012), but is also congruent with regional susceptibility in AD, where sensory cortices are relatively spared compared to the hippocampus. It is therefore possible that the weaker neurovascular function in the hippocampus could contribute to its earlier damage during AD.

Whilst the majority of AD cases are sporadic, the risk of going on to develop the disease is increased 9-15 fold by the possession of an ε4 allele of the apolipoprotein E (APOE) gene (Corder et al., 1993; Yamazaki et al., 2019). ApoE is a protein involved in the transport of lipids and is produced primarily by astrocytes in the central nervous system, but also by vascular mural cells and microglia, as well as by neurons under stress conditions, (Yamazaki et al., 2019). APOE4 expression exacerbates multiple features of AD, increasing Aβ accumulation and tau phosphorylation and altering synaptic function (Najm et al., 2019)(Yamazaki et al., 2019). It also decreases blood brain barrier (BBB) integrity and causes pericyte damage in animal models and human subjects (Bell et al., 2012); (Montagne et al., 2020);). Additionally, in anaesthetised mice, APOE4 mice had reduced cerebral blood flow (CBF) and smaller increases in CBF after sensory stimulation (Bell et al., 2012); (Koizumi et al., 2018).

Because in humans, CBF changes occur before any other AD pathology (Iturria-Medina et al., 2016), such APOE4-mediated vascular dysfunction could be an important contributor to the earlier emergence of AD in APOE carriers. However, gaps in the data remain, complicating understanding of how APOE4 genotype could be promoting AD.

Firstly, neuronal activity has largely not been studied concurrently with vascular function and reports as to the impact of APOE4 genotype on neuronal activity are variable. Large decreases in neuronal activity have been reported in mice aged only 4 months (Bell et al., 2012), but neuronal hyperactivity has been reported in old APOE4 mice (Nuriel et al., 2017). The potential direction of causality is therefore unclear: could decreases in CBF be due to decreased neuronal drive or could they themselves be restricting neuronal function?

Secondly, the impact of APOE4 on neurovascular coupling in the hippocampus has not yet been studied, despite it being more sensitive to AD damage (Henneman et al., 2009), a site of BBB breakdown in old APOE4 carriers (Montagne et al., 2020) and having weaker neurovascular function than more studied neocortical regions (Shaw et al., 2021a).

Finally, animal studies in which CBF and functional hyperaemia are dramatically decreased in young mice (Bell et al., 2012; Koizumi et al., 2018)do not recapitulate the human condition in which young APOE4 carriers show, if anything, increased CBF compared to APOE3 carriers from young into early old age (Thambisetty et al., 2010; Wierenga et al., 2013). BBB changes in human APOE4 carriers have also only been reported from old age (Halliday et al., 2016; Montagne et al., 2020), whereas these changes were observed in very young mice (Bell et al., 2012). It is therefore hard to interpret the relevance of rodent studies that show much more extreme changes in young animals (Bell et al., 2012; Koizumi et al., 2018) than are seen in humans.

An explanation for the greater impact of APOE4 in rodent studies than humans may be that carrying APOE4 increases the impact of stressors such as anaesthesia (Browndyke et al., 2021; Schenning et al., 2016). Inhalation anaesthetics such as isoflurane - the anaesthetic used in several of the rodent studies – seem to have a particularly large effect, reducing cognitive function acutely after surgery, before recovering to baseline levels after 10 days (Cai et al., 2012).

To plug these gaps in our understanding of how APOE4 impacts on cerebrovascular function, we therefore studied awake mice expressing human APOE3 or 4 in place of murine APOE, to avoid the confounding effect of acute surgery and anaesthesia. We did this by implanting mice with a chronic cranial window and recording neurovascular function up to 4 months after mice recovered from surgery, tracking function from young age to late middle age. To unpick the contribution of neurons and blood vessels to any alterations in neurovascular function we used 2-photon imaging to measure neuronal and vascular activity at a single vessel and neuronal level, as well as measuring the summed activity of neuronal metabolism and vasodilation using haemoglobin spectrometry and laser doppler flowmetry. Finally, we studied both the primary visual cortex (V1) and the CA1 subfield of the hippocampus to probe whether APOE4 effects were more pronounced in the hippocampus given its sensitivity to AD and weaker basal neurovascular function.

Instead of the dramatic reductions in CBF that were reported in acutely anaesthetised experiments, we observed several subtle yet persistent alterations in neurovascular function that persisted from young to late middle age, including impaired vasomotion, neuronal hyperactivity and less reliable vascular responses to neuronal activation. These data are more consistent with the pattern of results observed in humans than in anaesthetised mice, and suggest APOE4 genotype does not itself catastrophically impair vascular function but rather more subtly shifts neurovascular physiology such that it is more susceptible to further insults (e.g. surgery, ageing or beta amyloid accumulation).

## Results

### Reduced vascular density in young APOE4-TR mice is not reflected in baseline flow and blood oxygenation

Studies using anaesthetised preparations have previously shown decreased vascular density and cerebral blood flow in APOE4 compared to APOE3 TR mice (Bell et al., 2012); (Koizumi et al., 2018). We confirmed the reduction in vascular density in the visual cortex and HC of 3-4 month APOE4-TR mice using fixed brain tissue with the vasculature filled with a fluorescent gelatin (Fig. 1A, B). As previously (Shaw et al., 2021a), we also observed a reduced vascular density in HC compared to V1. The combined effect of the regional and genotype effects was that the vascular density in APOE4 mice HC was less than half that in APOE3 V1, suggesting potentially compromised function in APOE4 HC.

**Figure 1.**
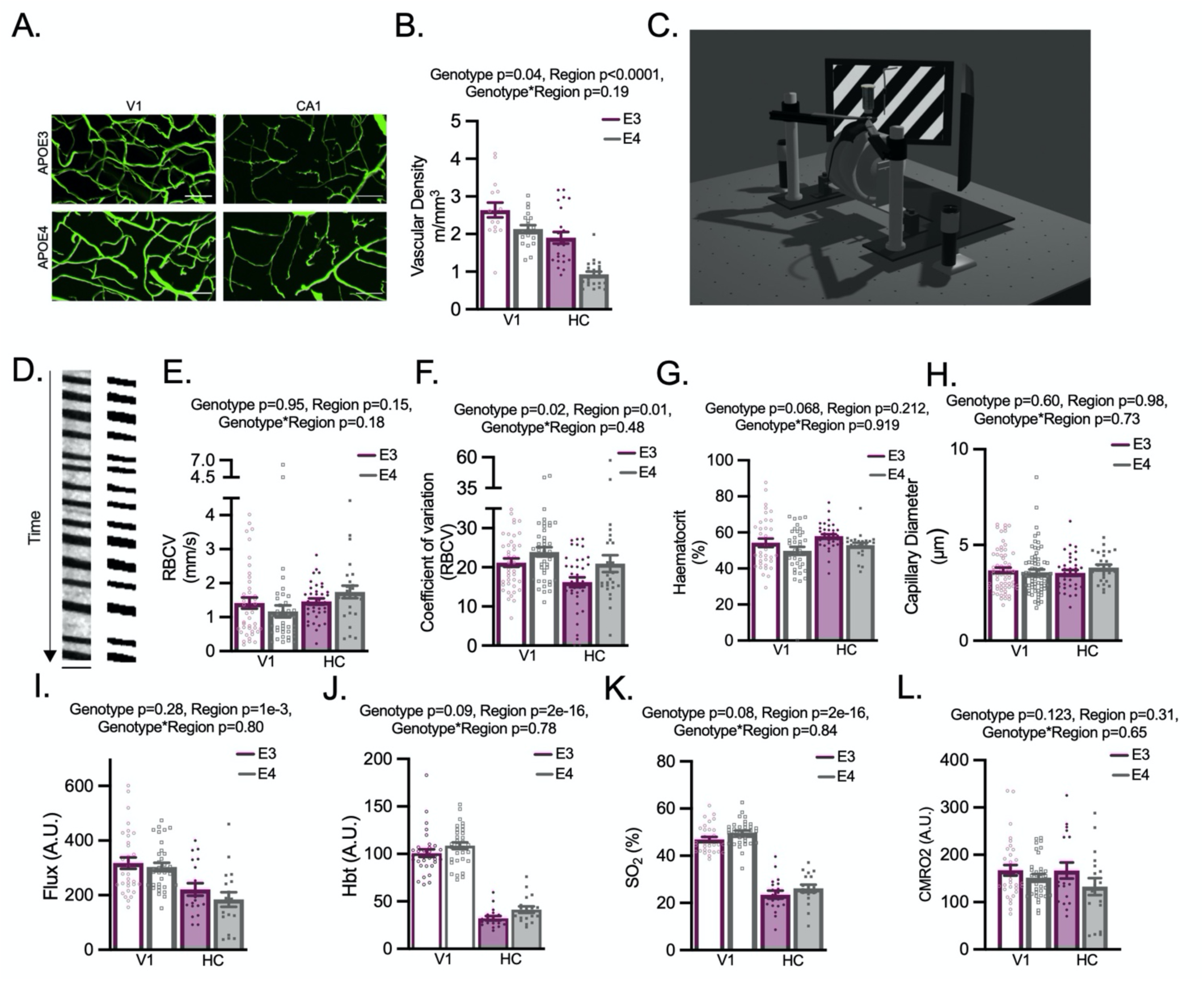
Under baseline conditions functional measurements are largely preserved in APOE4 mice within brain regions, but vascular density is reduced. **A)** Example z-projected images of FITC-gelatin perfused vessels (green). Scale bars: 50 μm. Projected depth: 45.8µm. **B)** Vascular density (of vessels <10 µm in diameter) was reduced in APOE4 animals in V1 and HC. Density was also significantly lower in HC than V1. **C)** Schematic of in vivo imaging set up. Screens display a visual stimulation (drifting grating) as presented to animals with a V1 surgery. Net haemodynamic measurements were recorded using the Oxy-CBF probe and individual vessels and neurons were imaged using 2 photon microscopy. **(D)** Example output from a capillary line scan after preprocessing (left) and the same image binarised (right) for haematocrit measurements. Scale bar (y) = 215ms, (x) = 4.8 µm. No region or genotype differences were found in RBCV **(E)** or in haematocrit **(G)** measures, but RBCV coefficient of variation (CV) was increased in APOE4 animals across regions, and decreased in HC compared to cortex **(F)**. **(H)** No genotype differences were observed in capillary diameters. Regional differences in net measurements of (I) flux, **(J)** HbT and **(K)** SO_2_ were observed, with values being lower in HC vs V1. Trend level increases were observed in HbT and SO2 measures of APOE4 mice. Individual dots on bar plots represent individual slices (B), individual vessels (E-H) and individual recording sessions (I-L). See appendix (i) for sample sizes and details on statistical tests. See also: Figure 1: Supplementary Figure 1

We then investigated the impact this reduction in vascular density had on various aspects of haemodynamic function, using awake head-fixed mice with either a cranial window implanted over the visual cortex or a cannula implanted over CA1 (Shaw et al., 2021a), at rest and in the absence of visual stimulation (hereafter referred to as ‘baseline’ conditions; Fig. 1C). Surprisingly, there were minimal differences in resting haemodynamics between genotypes, despite the large difference in vascular densities. Studying individual vessels using 2 photon microscopy, baseline red blood cell velocity (RBCV; Fig. 1D, E) and haematocrit (Fig. 1D, G) did not differ between genotypes or brain region, nor did the diameters of individual capillaries (Fig. 1H). The coefficient of variation (CV; calculated by dividing the standard deviation over time by the mean), reflecting temporal fluctuations in RBCV, was lower in HC than in V1 (Fig. 1F), suggesting more homogenous flow. This may promote oxygen extraction (Angleys et al., 2015) in hippocampal vessels, compensating for the lower hippocampal CBF. Conversely, flow was more variable in APOE4 than APOE3 vessels in both brain regions which may indicate less efficient oxygen extraction in APOE4 mice.

Macroscopic flow properties assessed using combined haemoglobin spectroscopy and laser doppler flowmetry also, however, revealed no differences between genotypes in RBC flux, total haemoglobin (HbT), oxygen saturation (SO_2_) or the cerebral metabolic rate of oxygen consumption (CMRO_2_) (Fig. 1I,J, K, L). Consistent with our previous findings (Shaw et al., 2021a), flux, HbT and SO_2_ were lower in HC than V1, while CMRO_2_ was the same. Therefore, the decrease in vascular density and increased coefficient of variation in APOE4 mice is not sufficient to drive net alterations in CBF or blood oxygenation, despite the absence of clear compensatory alterations in the diameter, RBC content or flow in individual vessels.

We then tested how much the measured vascular densities and blood SO_2_ levels might impact on tissue oxygen concentrations, using a simple model of oxygen diffusion into the tissue that we developed previously (Shaw et al., 2021a). Calculating blood oxygen concentrations from our measured SO_2_ values, we modelled tissue oxygen levels and oxygen consumption at different distances from blood vessels, based on measured vascular densities in each genotype. Our results indicate that both tissue oxygen concentration and oxygen consumption rates in brain tissue are no different between APOE genotypes in either brain region (Figure 1: Supplementary Figure 1).

### Neuronal activity is unchanged at baseline, but APOE4-TR animals display a hyperactive phenotype in response to visual stimulation

To allow us to relate vascular function to neuronal activity – i.e. to study neurovascular coupling – we bred APOE-TR mice with mice expressing GCaMP6f under the control of the Thy1 promoter, to generate mice which expressed GCaMP6f primarily in excitatory neurons (Dana et al., 2014) and were homozygous for APOE3 or APOE4 in place of murine APOE.

First, we characterised neuronal activity in the two genotypes. We imaged neuronal calcium activity in V1 and CA1 pyramidal cells (Fig. 2A) during baseline conditions, as well as evoked activity in the visual cortex during presentation of a drifting grating (Fig. 1C). We detected calcium events as peaks in the fluorescence signal in each region of interest (ROI) normalised to baseline (Fig. 2B). We assessed the number and size of individual calcium responses, how correlated cells’ firing was across the field of view (FOV) and measured the network response across all cells during visual stimulation. At baseline, neuronal activity was no different between APOE3 and APOE4 mice in either region (correlation, size or frequency of calcium peaks Fig, 2C-E)

**Figure 2.**
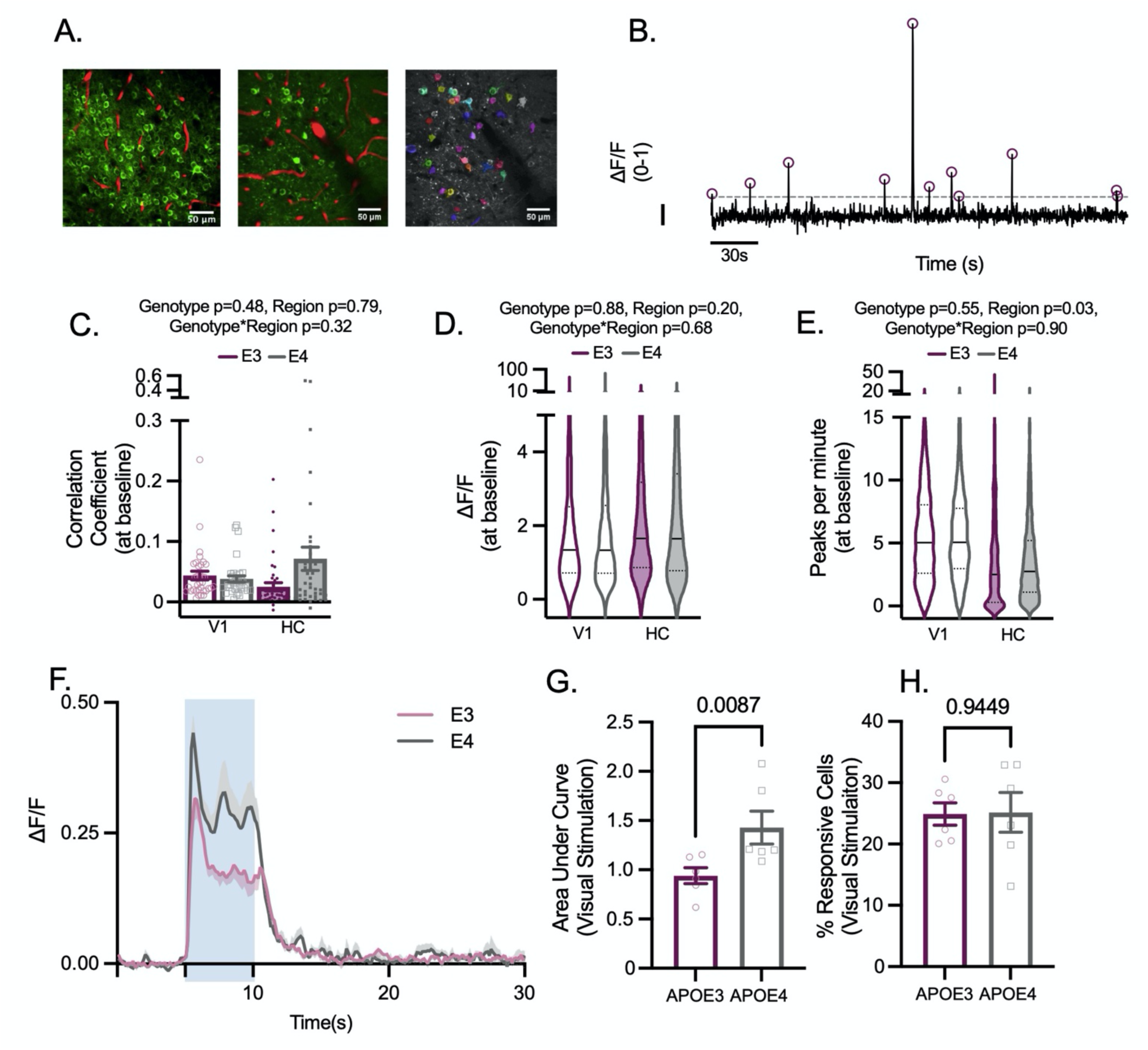
Under baseline conditions, genotypes display the same neuronal characteristics, but during visual stimulation APOE4 animals have larger neuronal responses. **(A)** Left: Thy1-GCaMP expressing neurons (green) and Texas Red filled vasculature (red) from CA1 (left panel) and visual cortex (middle panel). Right panel: ROIs overlying individual cells as detected using Suite2P (Pachitariu et al; 2016). **(B)** Example ΔF/F trace from one ROI, normalised between 0-1. Dashed line depicts threshold for peak selection, circles represent detected peaks and scale bar represents ΔF/F of 0.1 on the y axis and 30 s on the x axis. At baseline, **(C)** the correlation between individual cells within a recording, **(D)** the size of detected peaks per ROI, and **(E)** the number of detected peaks per ROI did not differ between genotypes. There were no regional differences in the correlation between cells or size of detected peaks, but the number of peaks per minute was lower in HC than V1. In visual cortex, the size of the neuronal response to 5s visual stimulation (blue shaded region) was larger in APOE4 animals **(F, G)** but the percentage of neurons that responded to visual stimulation did not differ between genotypes **(H)**. Individual data points plotted in barcharts represent individual fields of view (FOV, C) or animal averages **(G,H)** and violin plots represent values for individual cells. See appendix (i) for sample sizes and statistical tests.

In contrast, there was a larger change in fluorescence in response to a visual stimulation in APOE4-TR mice (Fig. 2F, G), though the number of cells that significantly increased their calcium during visual stimulation was unchanged (Fig. 2H). Therefore, calcium signals increase more in cells responsive to a visual stimulus in APOE4 compared to APOE3 TR mice. This effect could be due to increased calcium signals to the same level of electrical excitation or increased depolarisation or spiking of these cells during visual stimulation. Both options are plausible: calcium handling has been shown to be impaired in astrocytes of male APOE4-TR mice (Larramona-Arcas et al., 2020), while neurons of old APOE4-TR mice have been shown to be hyperactive due to decreased inhibitory GABAergic tone (Nuriel et al., 2017). We could not test whether a similar enhanced response to stimulation occurs in the HC as neurons are not activatable by a specific “stimulus” in the same way as in V1.

### Vascular responses to visual stimulation are unimpaired in the capillary bed, but pial arterioles are less responsive in neocortex APOE4-TR mice

Alterations in pericyte function have been previously reported to increase BBB permeability (Bell et al., 2012) and the decreased functional hyperaemia observed in APOE4 mice undergoing acute surgery suggests that the ability of vessels to respond to increased neuronal activity might be altered in APOE4 carriers (Koizumi et al., 2018). We used two separate approaches to study neurovascular coupling in HC and V1, because of different experimental constraints in the two regions. GCaMP6f expression was patchier in V1, so it was not always possible to record calcium signals next to imaged blood vessels, but we could record visual stimulus-evoked responses from the pial arterioles into the capillary bed. However, in HC, whilst we could not measure stimulus-evoked responses, and the large feeding vessels were too deep to be imaged, but we could image capillaries and small arterioles adjacent to spontaneous local neuronal calcium signals.

In V1, we measured vascular diameter changes in response to stimulation in pial arterioles and downstream capillaries, as well as stimulus-evoked changes in RBCV in capillaries. In a healthy system, visual stimulation evokes neuronal activity in the visual cortex and blood flow to the area increases accordingly, seen as increased dilations in capillaries and upstream pial vessels, and an increase in RBCV in individual vessels (Fig. 3E,A, I). The frequency and size of dilations in response to visual stimulation were the same in the capillary bed of APOE3 and APOE4-TR mice (Fig. 3F, G, H). However, the frequency of upstream pial dilations was decreased in APOE4-TR mice, as was the percentage of vessels in which RBCV increased during visual stimulation (Fig. 3D, L). Furthermore, the size of these responses, when they did occur, was no different between genotypes (Fig. 3B, C, J, K), suggesting the potential for dilation remained the same in both genotypes. This was also the case when the size of the pial arteriole response was normalised to the larger neuronal responses in APOE4 vs APOE3 TR mice to generate a neurovascular coupling index (NVCindex; Figure 3: Supplementary Figure 1). Because vasodilation propagates upstream from the capillary bed (Chen et al., 2014; Iadecola et al., 1997), this decreased reliability of APOE4-TR mice pial vessels in the absence of changes in capillary responsiveness may suggest an impairment in the propagation of dilation from the capillaries, rather than in the capacity of pericytes and smooth muscle cells to dilate.

**Figure 3.**
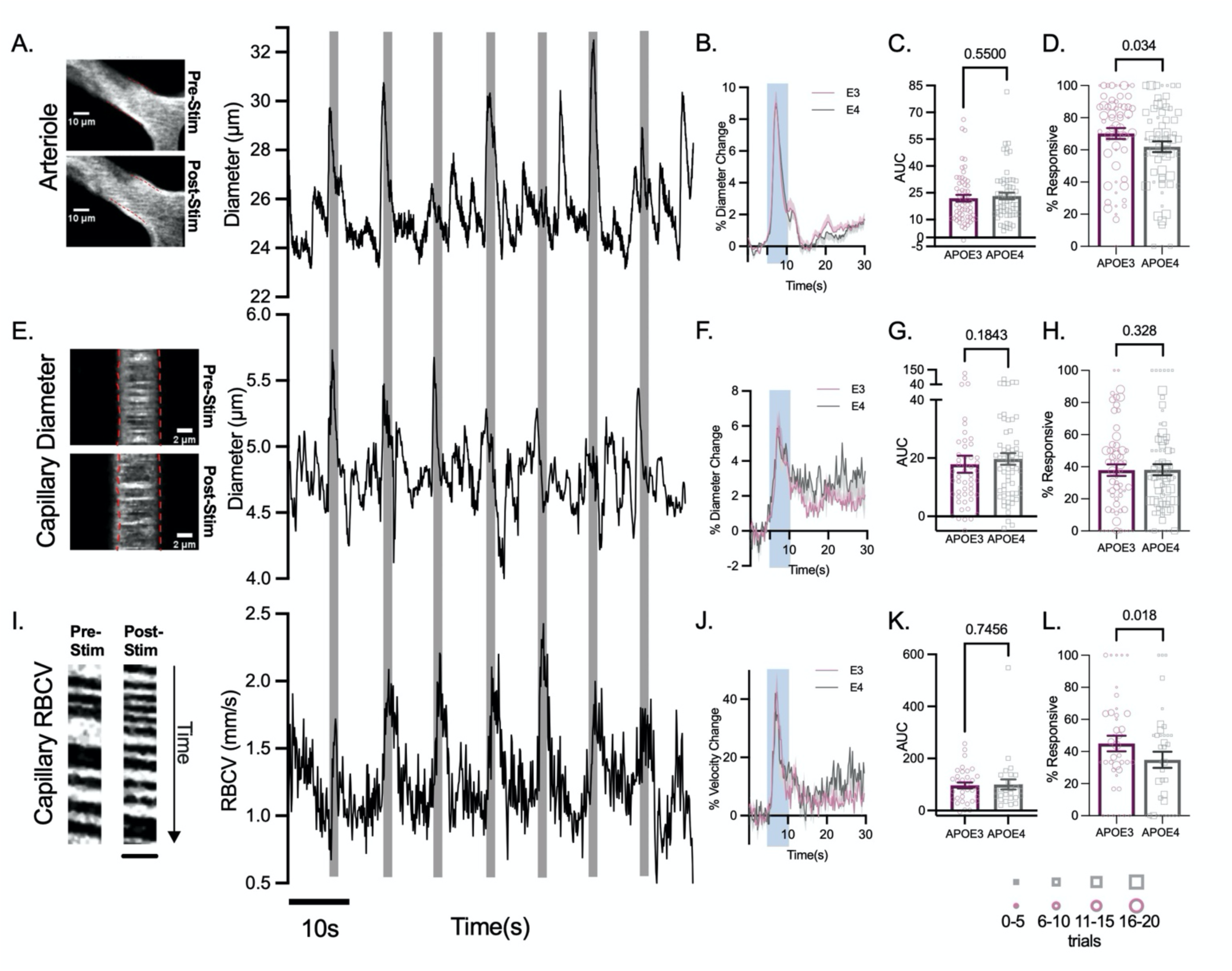
Vascular responses to visual stimulation are less reliable in visual cortex of APOE4 mice, but response sizes do not differ between genotypes. Example images and time series traces of pial arteriole diameter **(A)**, capillary diameter **(E)** and capillary RBCV **(I)** responses to visual stimulation (Scale bar (y) = 260ms, (x) = 5.6 μm). The response magnitude did not differ between genotypes in either the arteriole diameter **(B, C)**, capillary diameter **(F, G)** or capillary RBCV **(J, K)**. In APOE4 mice, the pial arteriole **(D)** and RBCV **(L)** response frequency was reduced, however there was no difference in the frequency of response to visual stimulation in capillary diameters **(H)**. Response frequency was weighted according to the number of contributing trials per vessel. The size of the individual data points reflects the number of trials, as per legend. Individual dots on bar plots represent the data points for individual vessels. See appendix (i) for sample sizes and statistical tests. See also Figure 3: Supplementary Figures 1 and 2.

However, these alterations in vascular responsivity did not affect the overall haemodynamic response, with CBF, sO_2_ and blood volume (total haemoglobin: HbT) responses to visual stimulation the same across genotypes (Figure 3: Supplementary Figure 2). The neurovascular changes observable at the single vessel level are therefore not, in these conditions, sufficient to affect these macroscopic measurements of functional hyperaemia. The lack of a genotype effect on overall CBF is consistent with the unimpaired capillary responses, as the capillary bed represents the majority of the resistance in the vascular bed, so is expected to mediate most of the increase in CBF during functional hyperaemia (Blinder et al., 2013; Hall et al., 2014), but is different from previous reports in anaesthetised mice showing a substantial reduction in functional hyperaemia to whisker stimulation (Koizumi et al., 2018).

In the HC, where we could not record from pial vessels but could directly relate local calcium changes to single vessel dilations, we found that while the average responsivity per vessel was not different across genotypes (Fig, 4A), there were more blood vessels that never responded to local calcium events (Fig. 4B), and fewer calcium events resulted in a dilation of local vessels in APOE4 vs APOE3 mice (Fig. 4C), despite similar increases in local neuronal activity (Fig. 4D, E). As in the visual cortex, dilations were not significantly different in size when they did occur (Fig. 4F-H). However, the reduced reliability of vascular responses was not sufficient to reduce the net regional responses to net changes in neuronal activity, as assessed by the size of fluctuations in total haemoglobin, blood flow or sO2 following fluctuations in CMRO2 (Figure 4: Supplementary Figure 1).

**Figure 4.**
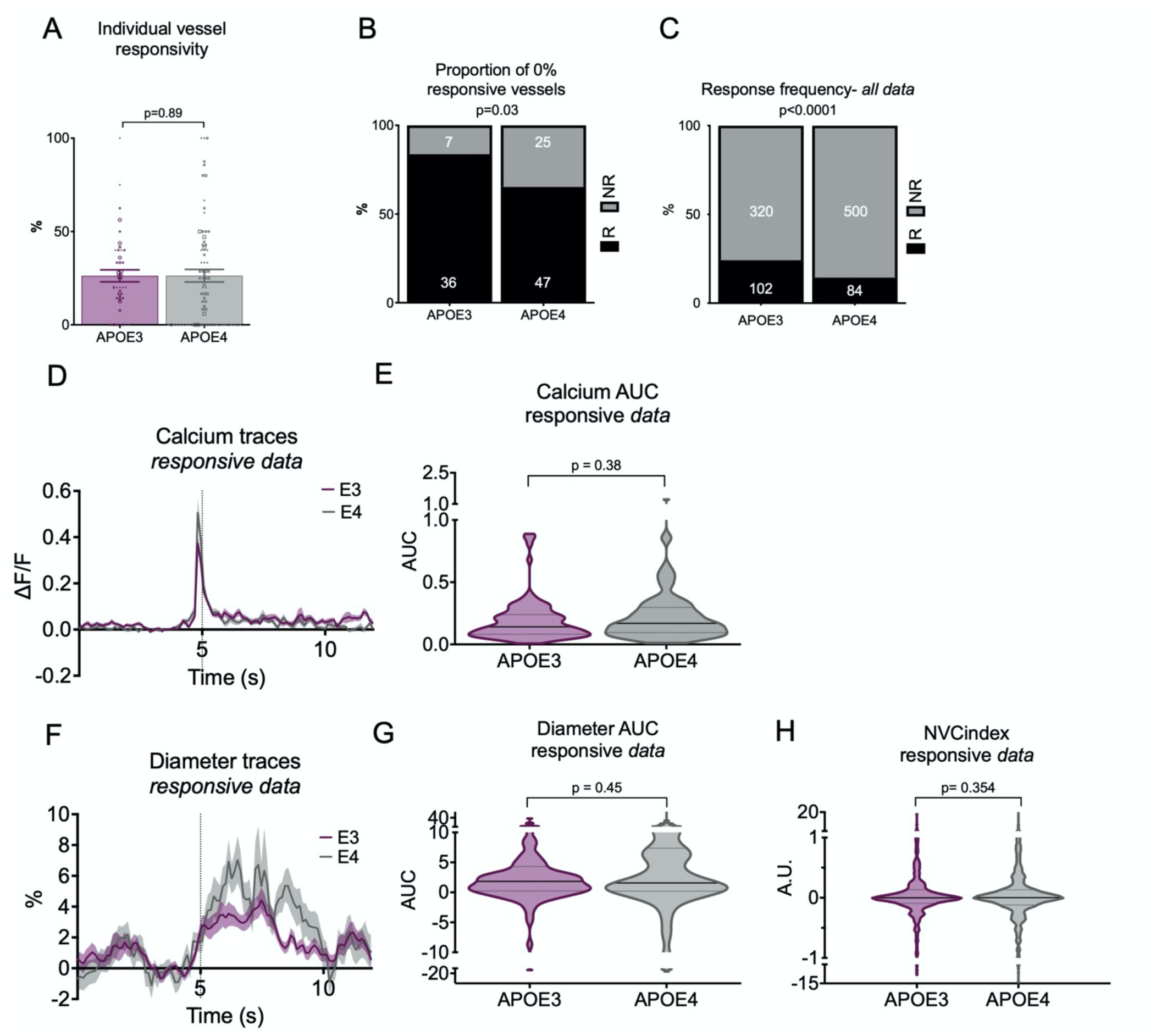
Vascular responses to preceding calcium events are less reliable in CA1 of APOE4 mice, but response sizes do not differ between genotypes. **(A)** Calcium evoked response frequency per individual vessel (for vessels with at least 3 contributing trials) was not different between APOE3 and APOE4 mice when data was weighted according to the number of contributing trials. The size of the individual data points reflects the number of trials, with the smallest dots corresponding with the data with the least trials (*symbol size 1: 3-5 trials, symbol size 2: 6-10 trials, symbol size 3: 11-15 trials, symbol size 4: 16+ trials*). **(B)** Local net calcium events were detected for each recorded vessel, and all recordings with >=3 calcium peaks were retained. Vessel responsiveness to the local net calcium event was classified, and overall response rates (%) were calculated per vessel. Significantly more vessels never responded (0% response rate) to preceding net calcium activity in APOE4 mice compared to APOE3 **(C)** Vessel response frequency to preceding individual calcium events was higher in APOE3 than APOE4 mice. **(D)** calcium peaks and **(F)** diameter traces were plotted for responsive dilations only (E, G) The AUC of the **(E)** preceding calcium peaks and vessel peaks **(G)** was not different between genotypes. **(H)** The neurovascular coupling index (dilation peak/calcium peak) was not different between genotypes. The individual values in the bar charts represent individual vessels and for D-H data points represent individual calcium events (and corresponding vascular responses). See appendix (i) for sample sizes and statistical tests. See also Figure 4: Supplementary Figure 1.

These results show that in both HC and V1, neurovascular coupling was mildly disrupted, as blood vessels dilated less frequently to increases in neuronal activity, but that these changes did not affect overall haemodynamics.

### APOE4-TR mice have impaired vasomotion

Arteries and arterioles oscillate at low frequencies (∼0.1Hz) — a phenomenon termed vasomotion (Mayhew et al., 1996). In addition to being indicative generally of vascular function and health (Das et al., 2021), vasomotion has also been shown to be one of the drivers of clearance from the brain (van Veluw et al., 2020) and is altered in mouse models of Alzheimer’s disease (Di Marco et al., 2015). As AD, and indeed APOE4, is associated with a reduction in clearance of Aβ, we tested whether vasomotion in pial arterioles in V1 (Fig. 5A) was affected by APOE genotype. Fourier transforms of our diameter traces (Fig. 5B) decompose the time course of diameter changes into a spectrum of the power of fluctuations at different frequencies, revealing peaks of increased power at the vasomotion frequency (∼0.1Hz) (Fig. 5C). However, the power at this frequency was strikingly lower in APOE4 mice (Fig. 5D), quantified by measuring the power at 0.1Hz (Fig. 5E). This was not because of differences in locomotion during recording (Fig. 5F).

**Figure 5.**
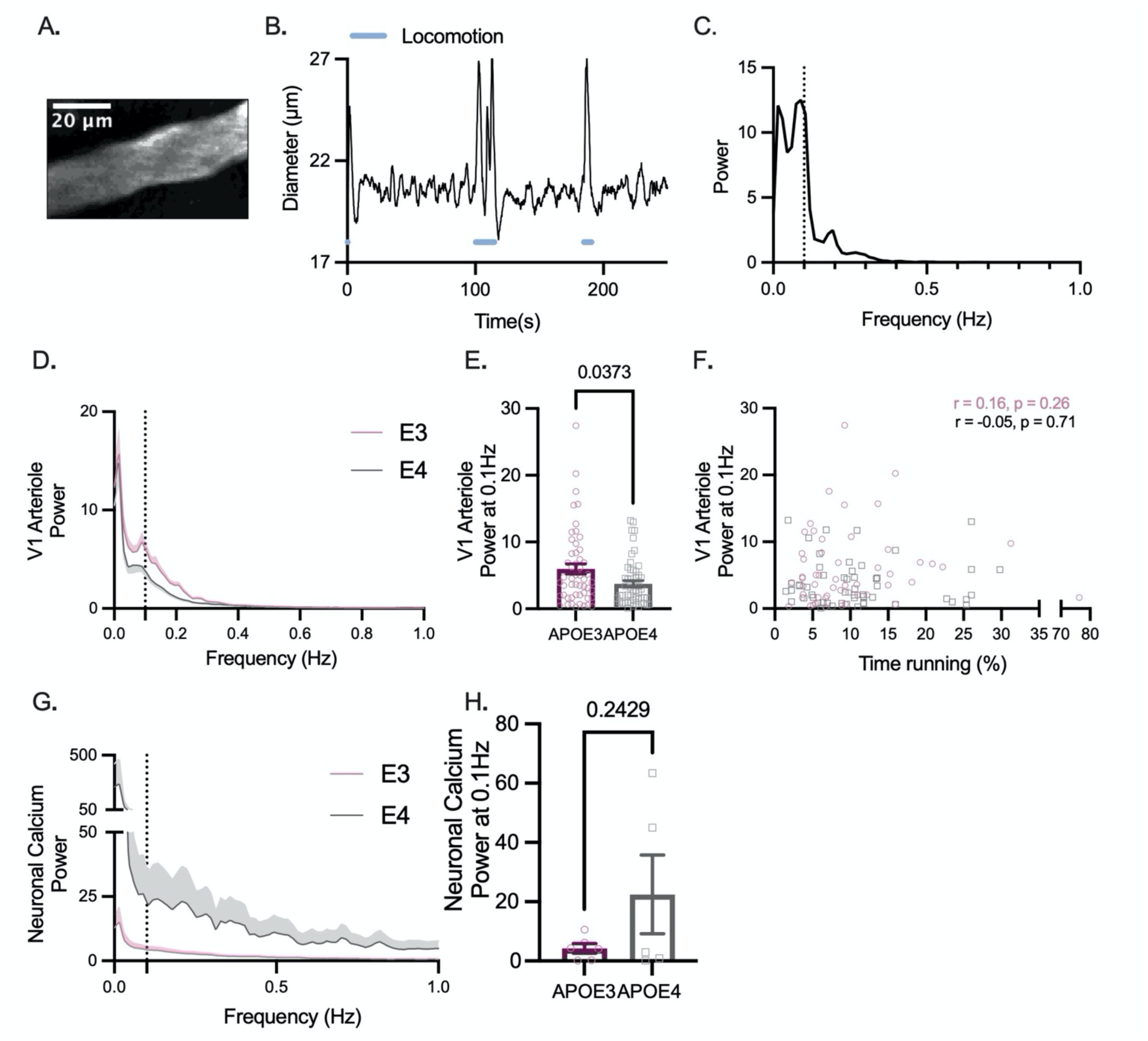
Vasomotion is reduced in the arterioles but not capillaries of APOE4 mice. **(A)** Example 2-photon image showing a representative pial arteriole from V1 **(B)** Example V1 pial arteriole diameter trace showing spontaneous fluctuations in diameter in the absence of locomotion (blue line) or sensory stimulation. **(C)** Power spectrum of diameter trace shown in B. Peak frequency can be observed at ∼0.1Hz (dashed line). **(D)** Average power spectra of arteriole diameter traces from APOE3 (pink) and APOE4 (grey) mice. Dashed line at 0.1Hz. **(E)** Arterioles from APOE4 mice have lower power at 0.1Hz. **(F)** The power at 0.1Hz was not affected by the amount of time spent running for either genotype. **(G)** Average power spectra for thy1-GCaMP6f fluorescence traces from V1 in APOE3 and APOE4 animals. (H) There was no observed difference in the power at 0.1Hz for neuronal calcium. Individual data points represent values for individual vessels (E-F) or animal averages**(H)**. See appendix (i) for sample sizes and statistical tests. See also Figure 5: Supplementary Figure 1.

It has been demonstrated by others that vasomotion, although an intrinsic vascular property, can be entrained by neuronal activity (Mateo et al., 2017), in particular the gamma band envelope, so we wondered if the decrease in vasomotion could be due altered neuronal activity across the frequency domain. Fourier analyses on GCaMP6f data, though acquired at too low a frequency to directly measure gamma band activity, can nevertheless reveal neuronal oscillatory activity that correlates with vasomotion (He et al., 2018). However, there was not only no peak in the neuronal power spectrum in the vasomotion range (Fig. 5G), but the power also did not differ between genotypes (Fig. 5H). That the APOE4-TR calcium power spectra, though very variable, were not reduced at low frequencies relative to the APOE3 TR signals, suggests that the observed reduction in oscillatory activity in APOE4 mice is likely vascular rather than neuronal in origin, at least as far as can be reflected in excitatory neuronal calcium signals. There were no observed peaks nor significant differences in power across genotypes at the vasomotion frequency in capillary diameters in V1 nor in vessel diameters recorded in CA1 (where we could not image the large feeding arterioles in HC that are equivalent to pial arterioles as these lie beyond our maximum imaging depth ; Figure 5: Supplementary Figure 1), consistent with our previous observations of low vasomotion in these vessels (Shaw et al., 2021b).

### Age does not modulate neurovascular alterations in APOE4 animals

For a subset of experiments in the visual cortex we were able to measure vascular and neuronal reactivity to visual stimulation at older ages, to investigate if increased age affected neurovascular function, as previous work has suggested a decrease in neuronal activity with increased age in APOE4 mice (Bell et al., 2012). Due to availability of appropriate GCaMP6f-positive mice, we could only measure neuronal activity at 3-4 and 6-7 months, while vascular features were also measured at 12-13 months. The size of the dilations of capillaries and arterioles decreased somewhat across genotypes between 3-4 and 6-7 months, but no consistent age-related decreases were observed up to 12 months (Fig. 6A-F). Where responses were affected by genotype in young animals, the same responses were generally seen in older animals, and no genotype effects emerged in any other metric. Specifically, the response frequency (Fig 6G) and vasomotion (Fig 7A-D) of pial arterioles remained significantly lower in APOE4 mice. Neuronal calcium during visual stimulation and baseline RBCV variability remained enhanced in APOE4-TR mice (Fig. 6J, K; (Figure 6: Supplementary Figure 1)). One measure - the frequency of RBCV increases to visual stimulation - was no longer different between APOE4 and APOE3 TR mice (Fig. 6I)) when studied across the whole age range studied, though not because of any significant differences between the ages studied.

**Figure 6.**
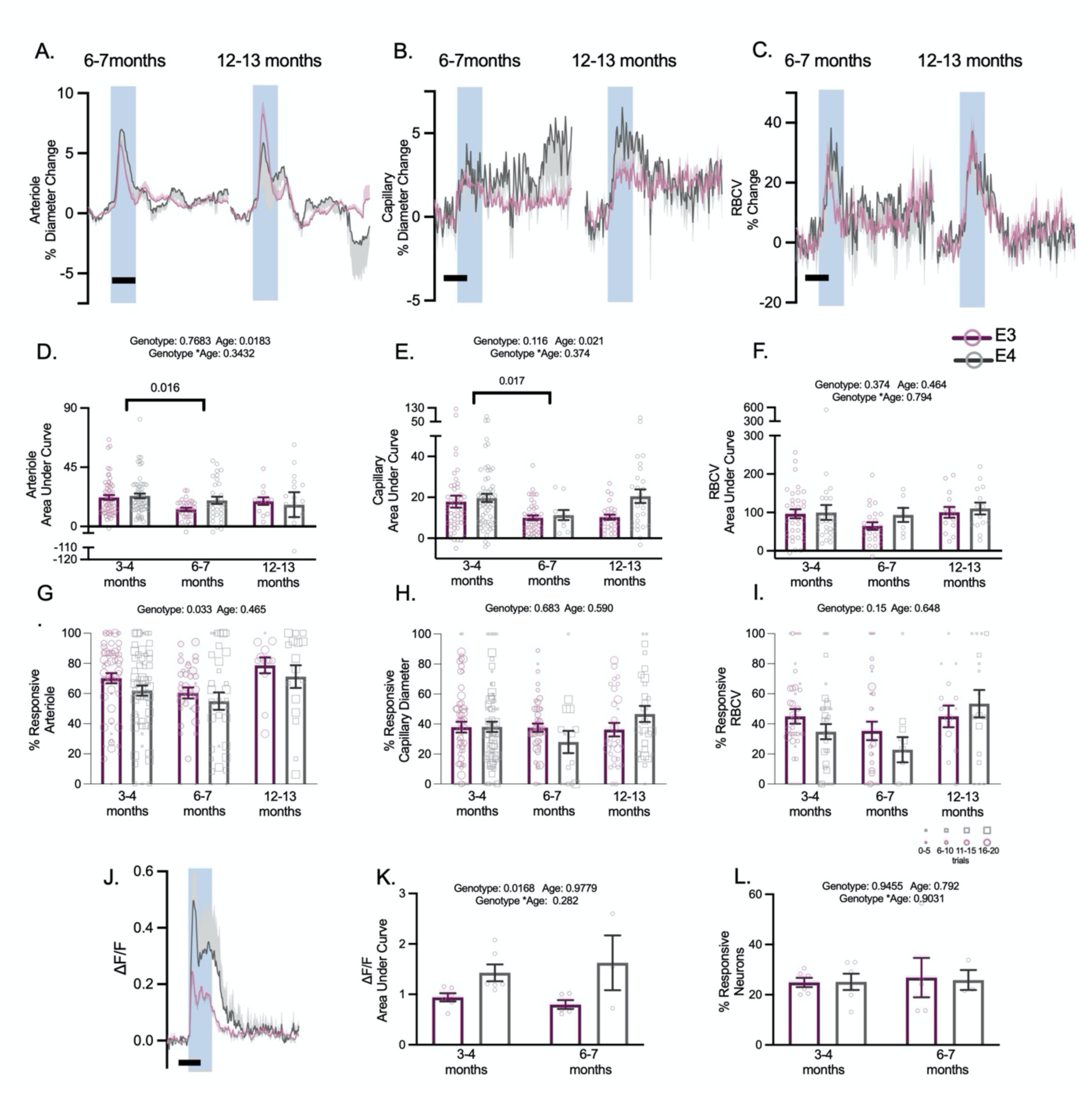
Age does not robustly modulate vascular or neuronal responses to visual stimulation. There was no genotype effect on the response magnitude to visual stimulation in animals across all age points **(A-F)**. In both arteriole diameter and capillary diameter measurements, there was an effect of age, with 3-4 month animals displaying larger responses to stimulation than at 6-7 months, however this effect was not observed in 12-13 month old animals. Response frequency to visual stimulation continued to be modulated by genotype across all ages in arteriole responses to stimulation **(G)**, however they were not affected by age. Neither capillary diameter response frequency **(H)** or RBCV response frequency **(I)** was modulated by age or genotype. At 6-7 months, neuronal responses to stimulation continued to be larger in APOE4 animals **(J, K)**, but this effect was not modulated by age. The response frequency of neurons to stimulation was not affected by age or genotype **(L)**. Individual dots on bar plots represent the values for single vessels (D-I) or animal average (K, L). Response frequency was weighted according to the number of contributing trials per vessel. The size of the individual data points reflects the number of trials, as per legend Scale bars = 5 s. Shaded regions represent the presentation of a visual stimulation. See appendix (i) for sample sizes and statistical tests. See also Figure 6:Supplementary Figure 1.

**Figure 7.**
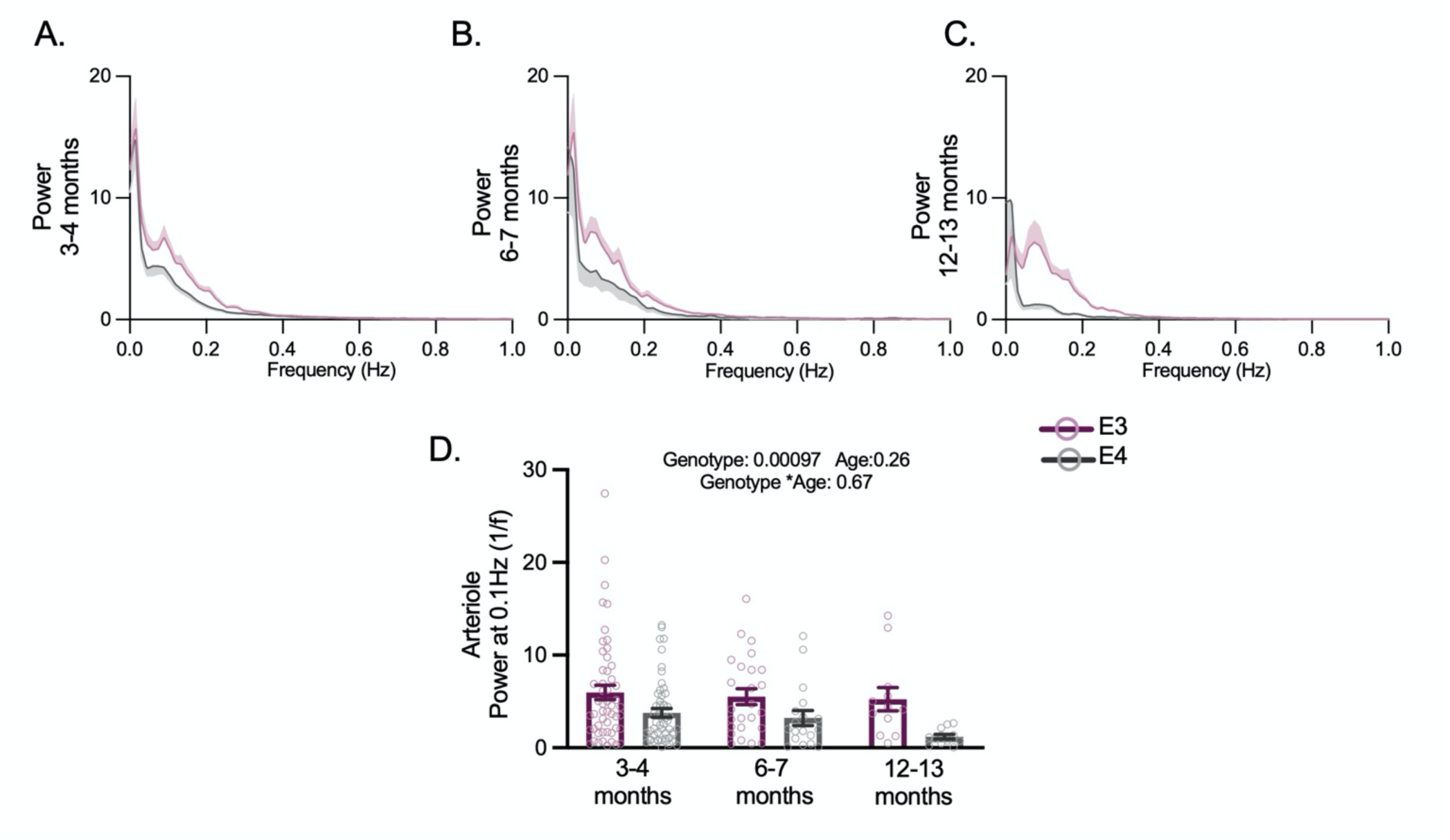
Vasomotion is reduced across all age points but does not appear to be modulated by age. Average power spectra of arteriole diameter traces across all three age points **(A, B, C)**. APOE4 animals had lower power at 0.1Hz **(D)**. Individual dots on bar plots represent the values for single vessels. See appendix (i) for sample sizes and statistical tests.

Together, these data suggest that, rather than declining with age, neurovascular function in APOE3 and APOE4-TR mice is stable, showing specific and persistent differences in pial arteriole responsivity, irregular vasomotion and neuronal function at least until late middle or early old age.

## Discussion

This was the first study to elucidate the effect of APOE genotype on neurovascular function at single neuron and vessel resolution in both the cortex and hippocampus of awake mice. Generally, we found milder effects of APOE4 genotype than has been previously suggested. While vascular density was reduced in APOE4 mice, this did not affect baseline CBF, blood sO_2_, or RBCV in single capillaries. Net increases in blood flow, sO_2_ or blood volume, or the size of single vessel dilations in response to visual stimulation were also not affected (unlike previously found for somatosensory stimulation (Koizumi et al., 2018). Pial and hippocampal arterioles were, however, less likely to dilate to neuronal activation, and in the cortex showed decreased vasomotion in APOE4 compared to APOE3 TR mice. Furthermore, unlike previously reported overall reductions in neuronal activity (Bell et al., 2012), we found an increase in neuronal calcium signals during visual stimulation. These genotype differences were unaffected by age, suggesting they represent a stable state that does not, itself, cause progressive dysfunction. These effects may, instead, interact with other factors to place the system more at risk of dysfunction when external events do occur (e.g., age, infection).

### Differences with previous studies

Previous studies have observed a large decrease in the CBF response to whisker stimulation in APOE4-TR mice (Koizumi et al., 2018), and a decrease in neuronal activity in response to hind limb stimulation, measured with voltage dyes (Bell et al., 2012). Although a regional difference between somatosensory and visual cortex cannot be ruled out, a more likely mechanism is the difference in experimental preparation. In our experiments, animals are imaged while awake and alert, having been allowed to recover from surgery for at least two weeks. The other studies utilised an acute preparation in which the mouse underwent cranial window surgery and was still anaesthetised when neuronal or haemodynamic recordings were undertaken. As human APOE4 carriers are more sensitive to anaesthesia than APOE3 carriers (Cai et al., 2012), mouse models of disease can be more susceptible to cortical spreading depression after invasive surgery (Shabir et al., 2020), and APOE4 is associated with an enhanced inflammatory phenotype (Kloske and Wilcock, 2020), the increased pathology observed in acute preparations may be due to this increased sensitivity to surgery and/or anaesthesia. Our findings, that there is an enhancement of neuronal activity and a slight disruption of vascular function, are a much closer approximation to the mild changes seen in young to middle-aged human APOE4 carriers (Henson et al., 2020; O’Donoghue et al., 2018).

### Physiological impact of APOE4 changes in neurovascular function

We initially hypothesised that the mild changes we observed in young animals might progress into worse pathology with age. We wondered if the increased neuronal activity observed in the visual cortex, suggesting an increased energy demand, might be insufficiently provided for by the less responsive vasculature leading to an energy imbalance that could eventually drive neuronal and vascular damage. Our data suggest that this is not the case. The alterations in V1 neuronal and vascular function remained largely stable from 3 to 12-13 months of age. Instead, therefore, our data suggest that APOE4-TR mice are different from APOE3 TR mice in specific, subtle, but consistent ways. This parallels findings in humans of the existence of subtle differences between healthy APOE3 and 4 carriers (higher default mode network functional connectivity in APOE4s), but the lack of age-dependent changes in any measures studied, from cognition to brain structure (Henson et al., 2020). The genotype differences do not, therefore, cause significant or progressive pathology in healthy, active individuals (our mice are housed in an enriched environment with access to an exercise wheel), but may cause APOE4 carriers (mice or humans) to be more sensitive to triggers that can then cause progression into a pathophysiological state. Such a trigger could be acute surgery under anaesthesia, as discussed above, experimental interventions such as decreased cerebral perfusion (Koizumi et al., 2018), environmental changes such as exposure to infection, a sedentary lifestyle, extreme age or, as in the two-hit hypothesis of AD, factors that initiate the accumulation of Aβ (Zlokovic, 2011).

A key issue for future research is then to untangle how existing differences between APOE3 and 4 carriers interact with additional triggers to cause a change from a stable state to progressive pathology and neurodegeneration. Because cardiovascular risk factors potentiate the risk of someone carrying an APOE4 allele developing AD (Juul Rasmussen et al., 2020), vascular factors are likely involved in this switch, while neuronal changes that could potentiate production of Aβ are also relevant. Some such APOE4-mediated differences are already known, including increased BBB permeability and pericyte inflammation (Bell et al., 2012; Montagne et al., 2020). Our results add three further features for consideration: increased neuronal calcium signals to stimulation, decreased vessel responsivity, and decreased pial vasomotion.

### Neuronal calcium

Before our study, increased neuronal calcium levels have been previously reported in cultured neurons exposed to ApoE4 both at rest and in response to stimulation (NMDA application, (Qiu et al., 2003); or mechanical injury, (Jiang et al., 2015); though in the latter, ApoE4 had no effect on resting calcium). Whether or not these observations and the increased signals we measure in vivo are due to increased neuronal depolarisation or altered calcium handling (as has been found in APOE4-positive astrocytes from male mice; (Larramona-Arcas et al., 2020)), the ubiquity of the involvement of calcium in cellular processes means that these alterations are likely to be physiologically relevant. The increase in calcium, and its ability to modulate synaptic strengths could therefore potentially underlie the increased connectivity and gamma band oscillations observed in human APOE4 carriers (Henson et al., 2020). It will be of interest to observe how such circuit changes develop in mice, to determine whether this contributes to the decrease in GABAergic inhibitory tone and electrophysiological hyperactivity that develops in old age APOE4 mice (Nuriel et al., 2017). Of particular interest is how these alterations affect how neurons respond to Aβ. Neuronal hyperactivity increases in response to Aβ, but predominantly in cells that are already active (Zott et al., 2019). Therefore, increased neuronal activity caused by ApoE4 may magnify the hyperactivity caused when Aβ is produced, exacerbating its pathophysiological effects.

### Vascular responsiveness

In our experiments, stimulus evoked pial dilations and calcium dependent dilation of CA1 arterioles and capillaries occur less frequently in APOE4-TR mice. These alterations in pial responsiveness were not sufficient to reduce the overall regional increase in CBF, HbT or oxygen delivery but do indicate something is altered in the pial vasculature of these APOE4-TR mice. Because the capillary bed responses were unchanged in the visual cortex, and the pial and CA1 dilations when they did occur were also of the same size in APOE3 and APOE4-TR mice, the capacity of the smooth muscle cells and pericytes of APOE4-TR mice to dilate may not be impaired, but instead the ability of dilations to spread upstream from the capillary bed to trigger upstream dilations may be reduced. A future research question would be whether endothelial calcium signals are altered as well as those in neurons, as these signals are involved in propagation of vasodilation via activation of transient receptor potential ankyrin 1 channels (TRPA1; (Thakore et al., 2021)) as well as in activating nitric oxide production, which itself can modulate propagation of vasodilation (Kovacs-Oller et al., 2020).

### Pial arteriole vasomotion

Possibly the most striking difference between APOE3 and APOE4-TR mice was the decrease in low frequency fluctuations of pial arteriole diameter in APOE4 mice - a phenomenon known as vasomotion. The mechanism underlying this reduced vasomotion is unclear, especially as we show that smooth muscle cells can dilate to neuronal activity with similar size responses to visual stimulation in APOE4 mice. Though vasomotion has been found to entrain to neuronal activity (Mateo et al., 2017), we find no evidence of a reduced neuronal drive on vasomotion, though this should be confirmed with electrophysiological measurements of local field potentials, to directly examine the gamma band envelope thought to entrain vasomotion.

Whatever the underlying mechanism, our results suggest another way in which, in APOE4 carriers, a system with reduced vasomotion could be stable, but at risk of decline if Aβ production increases: Vasomotion is known to be important for perivascular clearance of waste molecules, including Aβ, from the brain (Aldea et al., 2019) (van Veluw et al., 2020). APOE4 carriers have an increased Aβ burden both in AD and in cerebral amyloid angiopathy, probably as a result of reduced clearance (Greenberg et al., 2020). Whilst there are several mechanisms that have been shown to contribute to this, including reduced clearance across the BBB (Deane et al., 2008) and reduced degradation (Jiang et al., 2008; Miners et al., 2006), our data and others’ showing increased perivascular deposits of exogenously applied Aβ in APOE4 mice (Hawkes et al., 2012) point towards reduced perivascular clearance, due to a decreased driving force from vasomotion, as an additional mechanism. Thus, reduced vasomotion may not matter for the normal physiological functioning of the brain until Aβ production increases and clearance fails, allowing Aβ to accumulate further and exacerbate vascular dysfunction.

## Conclusion

Together our data point towards a subtle yet robust effect of APOE4 on the neurovascular system across the lifetime. They indicate APOE4-TR mice have increased neuronal calcium signals in response to sensory stimulation that may contribute to the altered network activity observed in humans and older mice, as well as specific deficits in vascular responsiveness and vasomotion that are not on their own sufficient to alter oxygen delivery or cerebral blood flow in healthy mice. However, the nature of these changes suggests that they may be important features that contribute to the decline of the previously stable system when an additional challenge is encountered, be that further vascular damage, infection or initiation of Aβ accumulation. Investigation of how such factors interact with APOE genotype should therefore be interrogated to understand how APOE4 genotype confers risk of developing AD.

## Materials and Methods

### Animals

All experimental procedures were approved by the UK Home Office, in accordance with the 1986 Animal (Scientific Procedures) Act, under a project licence held by C.H. All experiments were carried out on homozygous APOE3-TR or APOE4-TR mice (Knouff et al., 1999; Sullivan et al., 1997), crossed with animals expressing DsRed under the control of the NG2 promoter (Zhu et al., 2008) or with animals expressing GCaMP6f under the control of the Thy1 promoter. APOE-TR mice were bred from a colony at the University of Sussex derived from founders provided as a kind gift from N. Maeda (UNC School of Medicine, USA). All animals were on a C57BL/6 background. Both male and female mice were used in this study. Mice were between 3-4 months old unless otherwise specified. In ageing experiments, 6-7 month mice were the same as those imaged earlier at 3-4 months, while 12-13 month mice were a separate cohort. Animals were housed in a temperature-controlled room with a 12-hour light/dark cycle and had free access to food and water.

### Surgical Preparation

Animals were surgically implanted with a cranial window over the primary visual cortex or the CA1 subfield of the hippocampus as previously described (Shaw et al., 2021a). Briefly, animals were anaesthetised with isoflurane (∼4% at induction, ∼1.5 – 2% maintenance) and body temperature was maintained at 37°C using a homeothermic monitoring system (PhysioSuite, Kent Scientific Corporation). A 3 mm hole was created over the visual cortex or CA1, the dura removed, and for V1 surgery a glass window used to seal the exposed area. For CA1 surgery, ∼1.3mm of cortex was aspirated and a custom-made cannula with 3mm glass coverslip was inserted into the craniotomy and then secured in place at the skull surface. Custom-made titanium head bars were fixed to the exposed skull to enable later head fixation. Animals were administered subcutaneous injections of saline 0.9% (400μL), buprenorphine (1.2μg, Vetergesic, Ceva), meloxicam (6.2 μg, Metacam, Boehringer Ingelheim) and dexamethasone (120 μg, Dexadreson, MSD Animal Health) at induction, and continued to receive meloxicam (100 μg) in food for three days post-surgery.

### In vivo experiments

For all in vivo experiments, animals were head-fixed atop a cylindrical treadmill and locomotion was recorded using a rotary encoder (Kuebler). Recordings were made during both ‘baseline’ conditions, defined as the absence of visual stimulation with the animal free to engage in locomotion and, in animals with a window over V1, during visual stimulation. A drifting grating (315°, 2Hz full screen stimulus, with alternating spatial frequency trials of either 0.04 or 0.2 cycles per degree) was presented for 5 seconds on screens (Asus, ∼17cm from mouse), with an interstimulus interval of 20 seconds, allowing for stimulus triggered responses to be measured.

#### 2-photon microscopy

Animals were imaged using a 2-photon microscope (Scientifica) and a mode-locked Ti-sapphire laser (Coherent). Images were acquired using a 20x (XLUMPlanFL N, Olympus) or 16x (LWD, Nikon) water immersion objective from tissue excited at with a laser wavelength of 940nm or 970 nm. The objective was shielded with light occluding tape to minimise light artefacts from visual stimulation. Images were collected using SciScan (Scientifica) software.

Prior to imaging, animals were injected with a fluorescent dextran. Those with green fluorescence (i.e., GCaMP6f) were injected with 100µL 2.5% Texas red (Sigma-Aldrich) either subcutaneously (3 kDa) or intravenously via the tail vein (70 kDa), and those with red fluorescence (i.e., DsRed in vascular mural cells) were injected intravenously with 2.5% fluorescein isothiocyanate-dextran (70 kDa FITC-dextran; Sigma-Aldrich), allowing for visualisation of the vasculature. In <u>visual cortex</u>, pial arterioles were identified by the presence of smooth muscle cells (in NG2-DSred animals), or by their morphology and orientation relative to the large pial veins, as well as their response to visual stimulation. Pial arterioles were then followed with x-y recordings (average pixel size: 0.42 × 0.42 μm^2^, imaging speed range 3.8– 7.6Hz) made after bifurcations, until the vessels penetrated the parenchyma and therefore could no longer be classed as pial. Penetrating arterioles were followed and branching capillaries were imaged using high speed line scans, to record both diameter and RBCV (average pixel size: 0.19 μm, average number of traversals/second = 1224). In animals positive for GCaMP6f, additional x-y recordings (average pixel size: 1.28 × 1.28 μm^2^, imaging speed range 3.8–7.6Hz) were taken from the area proximate to vessel recordings at a depth that allowed for clear visualisation of neuronal cell bodies (layer 2/3 (∼z = -150μm)). In <u>CA1</u>, images were taken up to ∼500 μm from the surface of the window. Both x-y recordings of neuronal calcium across a large FOV (256 × 256 pixels, speed range 6.10–15.26 Hz, speed average 7.75 Hz, pixel size average 1.80 μm) and smaller FOV recordings allowing for concurrent vascular and local neuronal calcium signals (256 × 256 pixels, speed range 3.05–7.63 Hz, speed average 6.64 Hz; pixel size average 0.23 μm) were obtained. RBCV measurements were obtained as above using high speed linescans (average pixel size: 0.18 μm, average number of traversals/second = 791).

#### Oxy-CBF probe

Regional, or net (∼500 um^2^) haemodynamic measurements were recorded using a combined laser Doppler flowmetry/ haemoglobin spectroscopy probe (Oxy-CBF probe; Moor Instruments; (Royl et al., 2008) at an acquisition rate of 40 Hz. Net measures of total haemoglobin (HbT), blood flow (flux) and oxygen saturation (sO_2_) and cerebral metabolic rate of oxygen consumption (CMRO_2_ – calculated using equation (i) below) were obtained both at baseline and in V1, during visual stimulation.

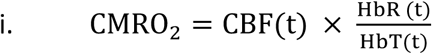

CBF represents cerebral blood flow, t represents time, HbR and HbT represents deoxygenated and total haemoglobin respectively.

### In vivo data analysis

#### Vascular diameter and RBCV measurements

Prior to analysis, several preprocessing steps were carried out in ImageJ (FIJI) to improve the quality of images as required. Such adjustments included registration, despeckling and/or median 3D filtering. In addition, all images had the ‘stack contrast adjustment’ plug-in applied to minimise any light artifacts arising from the visual stimulation. A custom MATLAB script was written to analyse vessel diameter using the full width at half maximum (Lab, 2021). Briefly, a skeleton was generated along the centre of each vessel and an intensity profile was plotted along a line perpendicular to the vessel. Values were obtained across a window of five vessel skeleton pixels and averaged to obtain a mean value per window. This was repeated at every second skeleton point, allowing for diameter measurements to be made along the full vascular arbour. These measurements were then averaged along the vessel, resulting in an average diameter per frame.

In high speed line scan experiments, similar methods were used to measure the diameter of capillaries imaged using windows of 40ms. Red blood cell velocity measurements were calculated with a radon transform using freely available code (Drew et al., 2010). In brief, the angle of shadows cast by the RBCs (that do not uptake fluorescent dextran) were measured and used to calculate the velocity across 40ms time windows that overlapped by 10 ms. Angles that were extremely large or small (because of motion artefacts) were removed. To further account for noise in the data obtained from line scans, traces went through either one (diameter) or two (RBCV) iteration(s) of outlier removal. In addition, for stimulation experiments, trials missing greater than 10% of data were excluded and those that remained were subsequently smoothed using a loess smoothing method (range: 1-5% span of the total number of data points, depending on which best represented the shape of the data). To determine AUC measurements, data was interpolated to fill in missing values, using a moving mean average.

A coefficient of variation was calculated for RBCV for each individual vessel, to measure temporal variation, by dividing the standard deviation over time by the mean value. To calculate haematocrit, images acquired of shadows cast by RBCs were binarised and the percentage area taken up by RBCs was calculated (black pixels vs. white pixels).

#### Neuronal fluorescence measurements

Regions of interest (ROIs) over individual neurons were identified in GCaMP6f recordings, using the freely available software Suite2p (Pachitariu et al., 2016). A fluorescence signal was extracted from each ROI, along with a background measure (F_neu_). To reduce out-of-focus neuropil contamination of the signal, 0.7 x F_neu_ was subtracted from each ROI (Chen et al., 2013).

#### Baseline analysis (no visual stimulation)

Neuronal measures: Baseline measures of neuronal activity were computed in rest (no locomotion) epochs of lasting at least 10 seconds by counting peaks per minute and the size of these peaks. To find peaks in the calcium signal, traces were first scaled so that all values fell between zero and one (equation ii). Peaks during rest periods were identified as being at least twice the standard deviation of the whole trace, and 0.25 seconds apart from the next peak. Each original trace was then normalised as is standard, (equation iii) and the size of each peak was determined and the number of peaks per minute was computed. During rest, the correlation of ROIs within a field of view was measured.

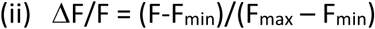

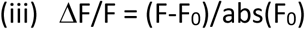

F represents the fluorescence trace. F represents the data trace (fluorescence), and F_0_ represents the baseline period to which the rest of the data is normalised.

Haemodynamic measures: For data that were recorded in the absence of a visual stimulation, rest periods were found as above for neuronal activity and an average value per vessel or animal (as specified in individual figure legends) was calculated for each measured parameter.

Detection of hippocampal vessel responses to local calcium events: Peaks in calcium traces were identified as being at least 1.5 times the standard deviation of the whole trace average, and 2 seconds apart from the nearest peak. The corresponding vessel traces (i.e. in the same FOV as neuronal region of interest) were then cut around the same time points. Responsive diameter traces were those where a dilation greater than 1 times the standard deviation of the baseline occurred for more that 0.5 seconds within 5 seconds of the calcium event. Response thresholds were chosen to best separate responses from non-responses.

#### Visual stimulation data analysis

Neuronal and vessel diameter data acquired from visual stimulation experiments in V1 were cut into trials around the stimulus presentations and averaged to yield a mean response per vessel or animal, as specified in figure legends. Only trials where there was no significant locomotion during the period two seconds prior to stimulation onset or during the stimulation period were used. Significant locomotion was defined as an event that was more than one third of a second in length and/or less than one second apart from adjacent locomotion epochs.

Data were normalised to the 5 s baseline preceding the onset of visual stimulation using equation (iii). To determine if there was a response to stimulation, a threshold was set as twice the standard deviation of this 5 s baseline period. Any response larger than this threshold was deemed ‘responsive’. For neuronal data all trials were averaged per cell and if the mean response was larger than the threshold, it was deemed a responsive ROI and subsequently averaged to provide a mean response size per animal. A percentage response rate was also calculated per vessel or animal. For vascular data, these trials were averaged per vessel to provide a ‘responsive only’ trace. Each vessel had a different number of contributing trials (because mice ran different amounts for each recording) and those with a low number of trials were likely to provide more extreme response frequency values (e.g., 0% or 100%). Therefore, the number of contributing trials per vessel was used to weight data when calculating response frequency. When calculating the size of responses, area under the curve (AUC) measurements were taken during the stimulation period using the inbuilt MATLAB function “trapz” which utilizes trapezoidal numerical integration. In. V1, neurovascular coupling indices (NVCindexndex) were calculated by dividing each vascular AUC by the average neuronal AUC for each genotype (as some mice did not express GCaMP6f in neurons, so calcium data was not available for each mouse). In CA1, where neuronal activity was measured local to the vasculature, the NVCindex was calculated by dividing responding vessel diameter peaks by the corresponding neuronal calcium peaks. Similarly in CA1, net haemodynamic peaks were divided by the corresponding CMRO_2_ peaks

#### Power spectrum analysis

Welch’s power spectral density estimates were computed across all traces (including locomotion epochs) for arteriole diameters, calcium fluorescence traces and capillary diameter traces. All spectra were computed from data recorded in the absence of visual stimulation (except for capillary traces in V1 which were collected during visual stimulation, however the expected increase in power as a result is not expected to be near 0.1Hz). All data was detrended by subtracting the baseline (the 8th percentile calculated over 15 second time windows (Grijseels et al., 2021)). Discrete Fourier transforms were then carried out across 60 second time windows using the inbuilt MATLAB function “pwelch”. Data below 1 Hz was selected and interpolated to the same length using linear interpolation. All data underwent outlier removal based on the value at 0.1Hz (all values greater than 3 standard deviations above the mean were removed). This resulted in the removal of a total of three pial arteriole traces (1 x APOE3, 2 x APOE4), two capillary traces in V1 (1 x APOE3 and 1x APOE4), three hippocampal vessel traces (1xAPOE3 and 2 x APOE4), and no calcium traces.

### Vascular density calculations

APOE-TR mice underwent transcardial perfusion under terminal anaesthesia. Animals were first perfused with 0.1 M phosphate buffered saline (PBS) then 4% paraformaldehyde (PFA) in PBS, and were then perfused with 5% gelatin containing 0.2% FITC-conjugated albumin (at 37 °C). Following at least 30 minutes on ice, brain tissue was then extracted and stored in 4% PFA at 4°C for 24 hours before being transferred to 30% sucrose in PBS for at least 3 days. Tissue was then sliced at 200µm on a vibratome, and slices were imaged using confocal microscopy (Leica SP8, pixel size: 0.45 – 0.57µm). Vascular density was calculated from the length of vascular skeleton per unit volume, using the “Analyse Skeleton” plugin in FIJI/ImageJ (Arganda-Carreras et al., 2010).

### Oxygen Modelling

Oxygen modelling was carried out as described previously (Shaw et al., 2021a). Briefly, median and upper lower 95^th^ centile sO2 measurements were used to calculate the pO2 in red blood cells, using the Hill constant for oxygen dissociation from haemoglobin in C57/BL6 cells (Uchida et al., 1998). The plasma pO2 was then estimated from comparison with similar measurements of intra- and interRBC pO2 to determine the capillary pO2(Lyons et al., 2016; Parpaleix et al., 2013). Distance maps were then calculated of 3D stacks of the vasculature of APOE3 and 4 visual cortex and hippocampus, coding every voxel in terms of the distance to the nearest vascular voxel. The median and 95^th^ centile spacing of voxels were calculated from the distribution of these values. Oxygen diffusion across these distances was then modelled by numerically solving Fick’s diffusion equation (iv) for radial geometries using the Partial Differential Equation toolbox in MATLAB, with a diffusion coefficient (D) of 9.24 × 10^−8^ m^2^/min in brain tissue at 37 °C (Ganfield et al., 1970)and 40% lower in a 200 nm vessel wall surrounding the capillary(Thomsen et al., 2017). Oxygen consumption by the tissue was assumed to follow Michaelis-Menten kinetics with a K_m_ of 1 µM, from the EC50 for oxygen at cytochrome c oxidase (Cooper, 2003), and a V_max_ of 2 mM/min (Gould et al., 2017).

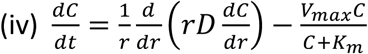

## Supporting information

Appendix containing full statistics reports for all analyses

## Statistics

Statistical analyses were conducted in GraphPad Prism 8, SPSS, RStudio or MATLAB. Data are presented as mean +/-SEM except for violin plots where horizonal lines represent the median and interquartile range. Individual data points represent individual blood vessels, brain slices, imaging sessions or animals as specified in figure legends. Where relevant, data whose residuals were normally distributed, as determined by a D’Agostino-Pearson and Shapiro-Wilk test, were compared using an unpaired t test, with a Welch’s correction applied if variances were unequal as determined by an F test. Data that were not normally distributed were compared using a non-parametric Mann-Whitney U test. Linear mixed models (LMM) were carried out using ‘lmer’ in RStudio, with random effects specified in appendix (i). For correlation analyses non-normal data were analysed using Spearman’s rank correlation. To weight response frequency data, so that vessels with a larger number of contributing trials contributed more to the mean, a weighted least squares linear regression was used and the mean was weighted by the number of contributing trials. P values below 0.05 were considered significant and those below 0.1 as trending towards significance.

## Funding

This work was funded by an Alzheimer’s Society DTC PhD studentship to O.B., Sussex Neuroscience PhD studentships for D.G. and D.C., the MRC (MR/S026495/1; MC_PC_15071) and an Academy of Medical Sciences/Wellcome Trust Springboard award to C.N.H.

## Acknowledgements

We would like to thank Jason Berwick and Jimena Berni for helpful discussions on this work.

## Author Contributions

O.B., K.S. and L.B. collected data for the studies. O.B., K.S., S.K. and C.N.H. designed the studies, and O.B., K.S., S.A. and C.N.H. analysed the studies. O.B, K.S, D.M.G, D.C and C.N.H. wrote scripts to analyse the data. O.B. and C.N.H. wrote the manuscript with feedback from all authors.

## Supplementary Information

**Figure 1: Supplementary Figure 1.**
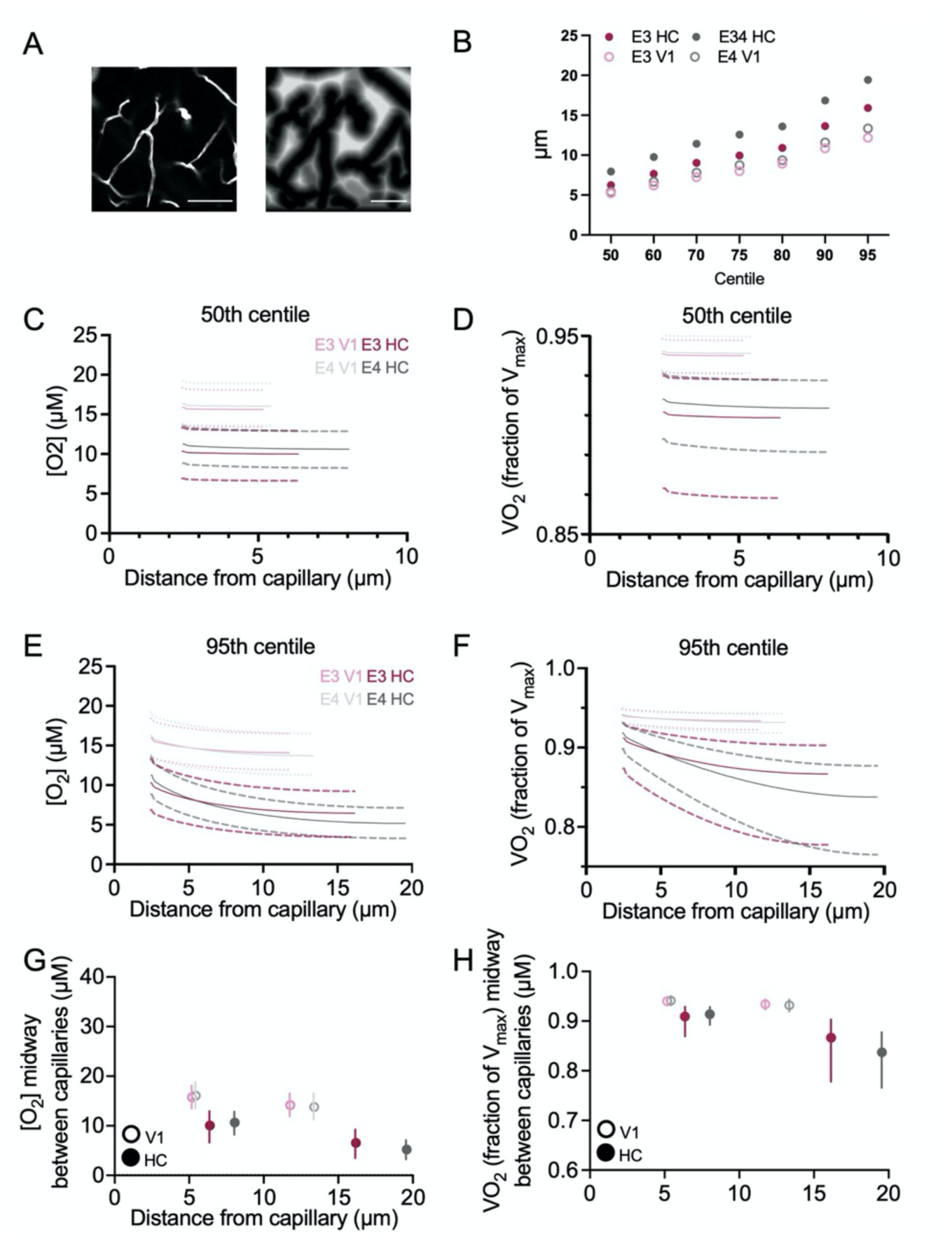
Modelling O2 diffusion in brain tissue. ***(A)*** Distance maps of ***c***onfocal z-stacks of FITC-labelled vasculature from APOE3 or 4 visual cortex and hippocampus were used to calculate the distance of each parenchymal voxel from a vessel in each condition. Scale bar: 40µm. **(B)** The median distance of a parenchymal voxel and increasing centiles of the distribution were used in the diffusion model. Oxygen gradients across the **(C)** median and E) 95% centile distances were calculated using a simple model of oxygen diffusion and consumption, as described previously (Shaw et al., 2021a). Oxygen concentrations at the vessel were calculated from the median (solid lines) and upper and lower 95 % confidence intervals (dashed lines) for SO2 measurements in APOE3 and APOE4 visual cortex and hippocampus, as described (Shaw et al., 2021a). **(D) & (F)** The rate of oxygen consumption in the tissue given these oxygen gradients was then calculated (VO_2_, expressed as a fraction of the rate if oxygen levels were not limiting for ATP production), using the known oxygen-dependence of oxygen consumption by cytochrome c oxidase, as previously. Finally, the oxygen concentration **(G)** and VO_2_ **(H)** for voxels at the median distance and 95% centile from a vessel are summarised. Error bars indicate models run with the 95% confidence intervals for SO_2_ measurements in the different conditions. Despite the lower vascular density in APOE4 mice, when taking into account the measured blood oxygen levels, oxygen and VO_2_ values are not predicted to be different in APOE3 and APOE4 mice in either the visual cortex or the hippocampus, based on the large overlap between genotypes of the 95% confidence intervals (Cumming and Finch, 2005), though as we found previously, both tissue oxygen levels and VO_2_ values drop significantly lower in the hippocampus than the visual cortex. Open circles represent values from the visual cortex, whilst filled circles represent values from the hippocampus. Light pink and light grey lines represent values from the visual cortex and hippocampus of E3 mice respectively. Dark pink and dark grey lines represent values from the visual cortex and hippocampus of E4 mice respectively.

**Figure 3: Supplementary Figure 1.**
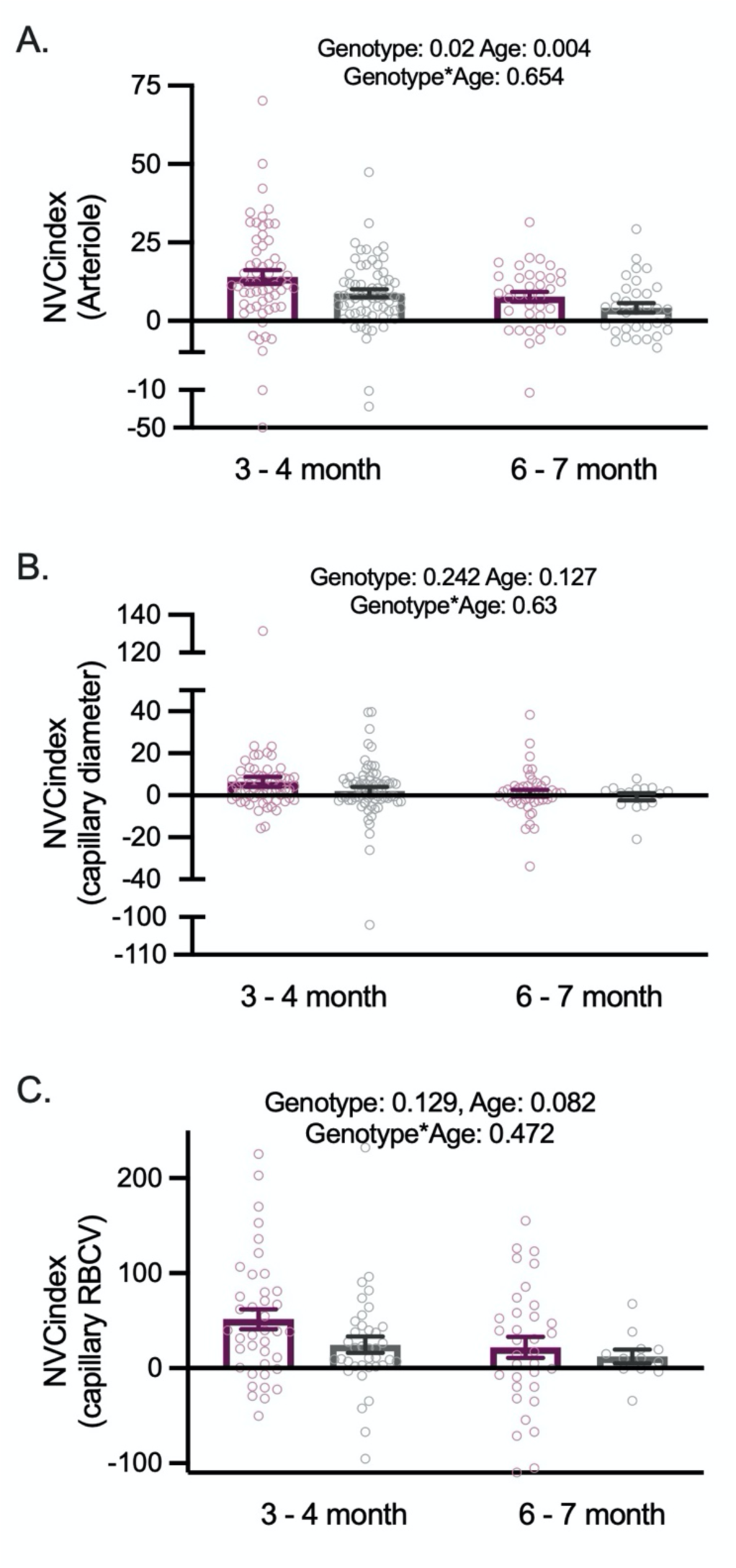
Vascular response to visual stimulation normalised to average neuronal response to produce a neurovascular coupling index (NVCindexndex). Raised NVCindexndex in APOE3 animals was observed in pial arteriole **(A)** responses to visual stimulation. No difference was seen in capillary diameter **(B)** or RBCV NVCindex **(C)**. Individual data points represent values from individual vessels. See appendix (i) for sample sizes and statistical tests.

**Figure 3: Supplementary figure 2.**
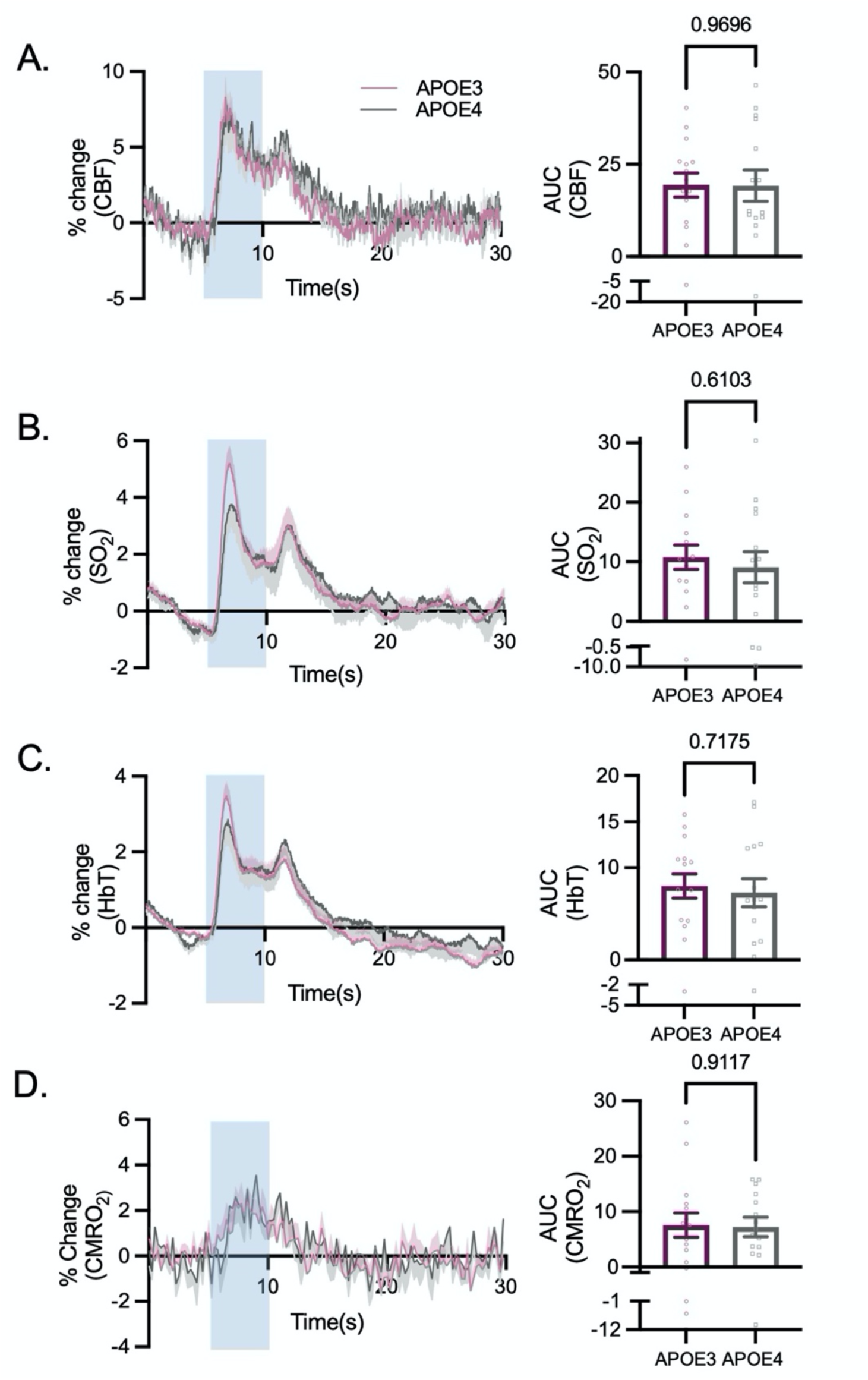
No effect of genotype was observed in net stimulus-dependent responses: Left panel shows average changes in net CBF **(A)**, sO_2_ **(B)** total haemoglobin (HbT) **C)** and cerebral metabolice rate of oxygen consumption (CMRO_2_); **D)** during 5s visual stimulation (blue shaded region). Right panel shows AUC for each during visual stimulation. Individual data points represent animal averages. See appendix (i) for sample sizes and statistical tests

**Figure 4: Supplementary Figure 1:**
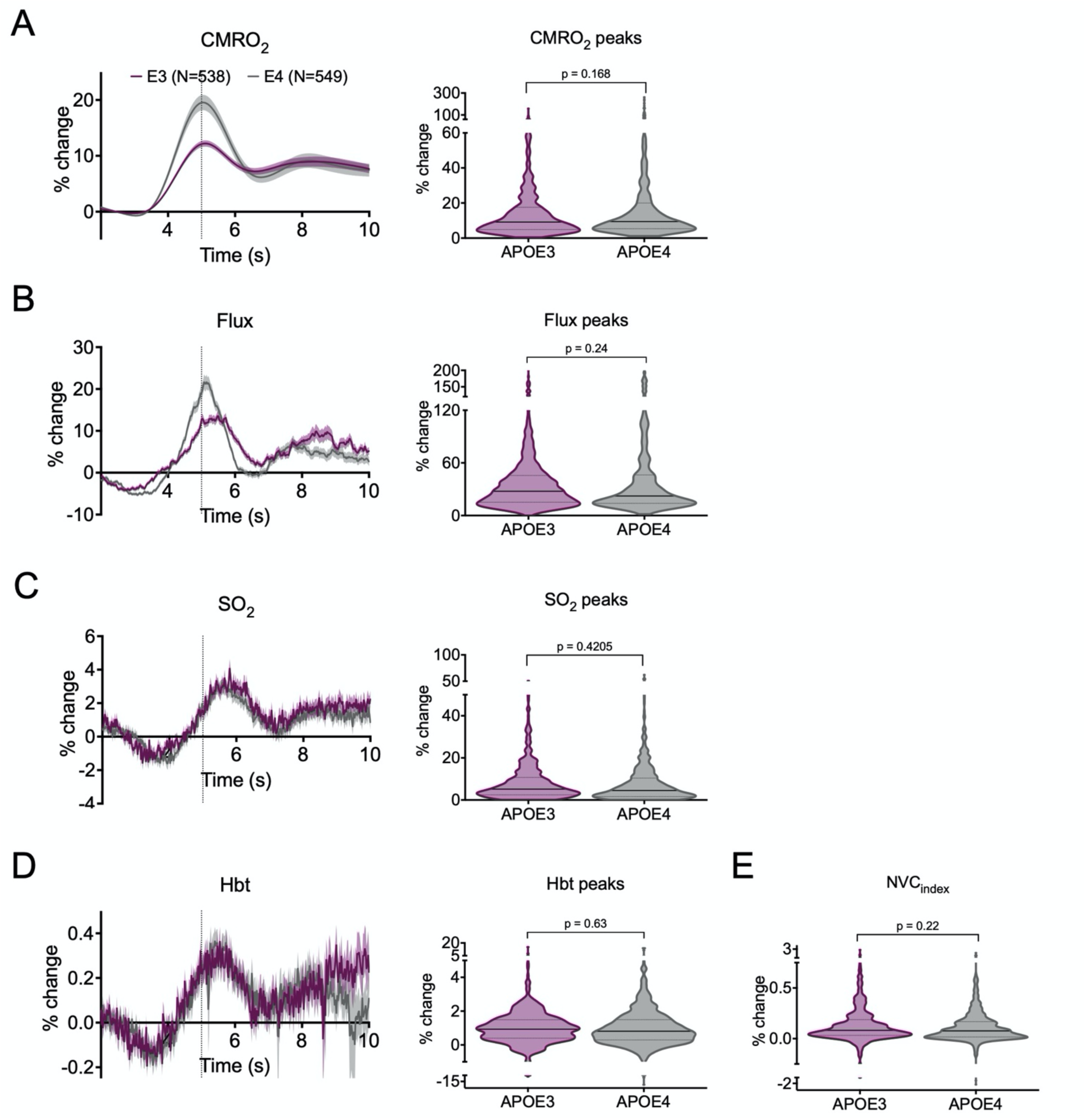
CMRO2-dependent haemodynamic responses in CA1. **(A)** Detected peaks in CA1 Oxy-CBF probe recordings of net CMRO_2_ were not larger in APOE4 than APOE3 mice, Corresponding CMRO2-dependent haemodynamic traces were then visualised for **(B)** flux, **(C)** oxygen saturation (SO_2_) and **(D)** total haemoglobin (HbT). There were no significant genotype differences in CMRO2-dependent flux, SO_2_ or HbT, **(E)** The NVCindexndex was calculated by dividing HbT/CMRO_2_, and showed no effect of genotype. Violin plots are composed of data points from individual haemodynamic events. See appendix (i) for sample sizes and statistical tests.

**Figure 5: Supplementary Figure 1:**
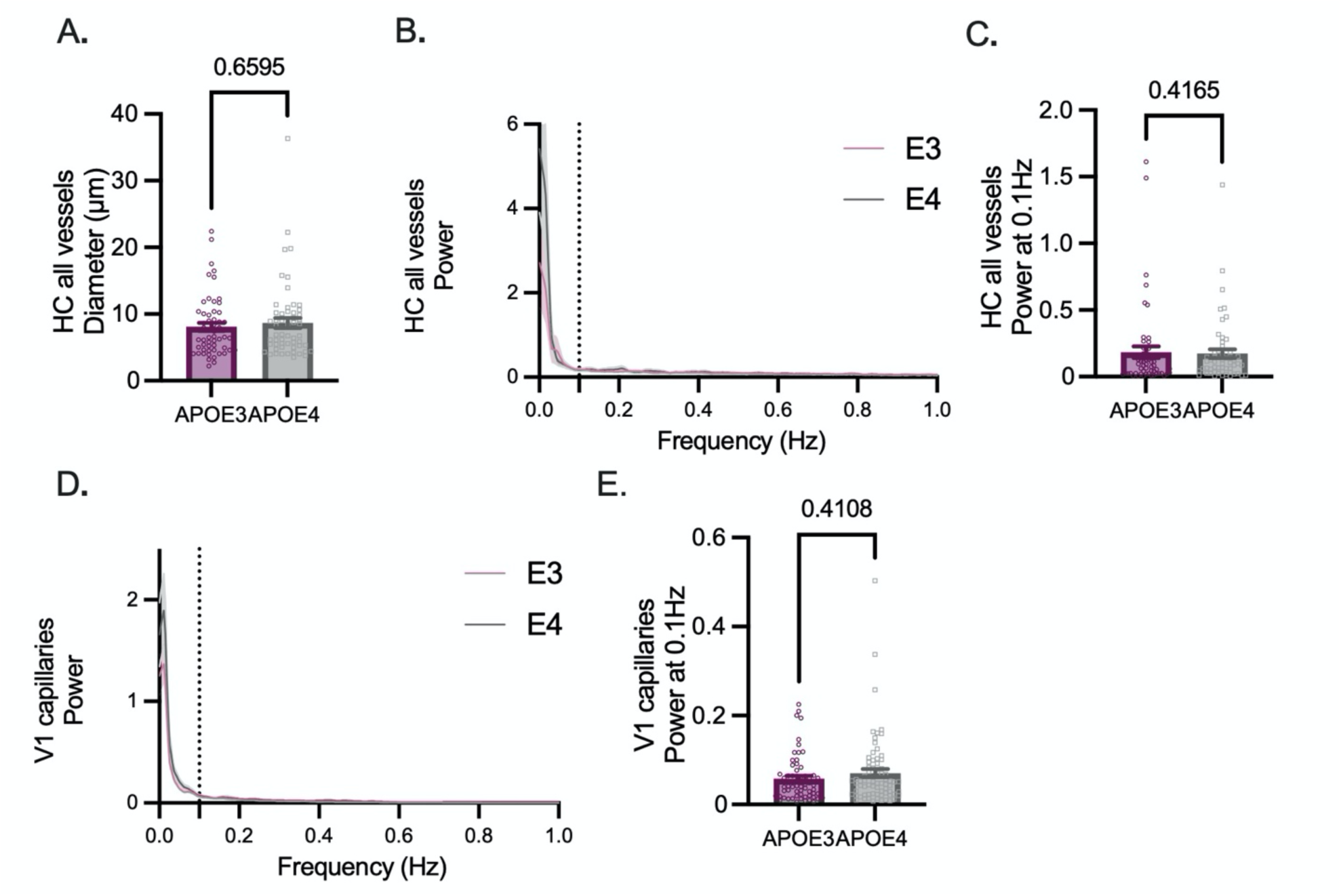
In APOE3 and APOE4 mice, power at the vasomotion frequency is the same in the vessels of the hippocampus and in the capillaries in V1. The diameter of **(A)** all the vessels recorded in CA1 were not significantly different between genotypes. **(B)** Average power spectra of CA1 diameter traces from APOE3 (pink) and APOE4 (grey) mice for all recorded vessels. Dashed line at 0.1Hz. **(C)** Power at 0.1Hz did not differ between genotypes. **(D)** Average power spectra of V1 capillary diameter traces from APOE3 (pink) and APOE4 (grey) mice. **(E)** Power at 0.1Hz did not differ by genotype. See appendix (i) for sample sizes and statistical tests.

**Figure 6: Supplementary Figure 1:**
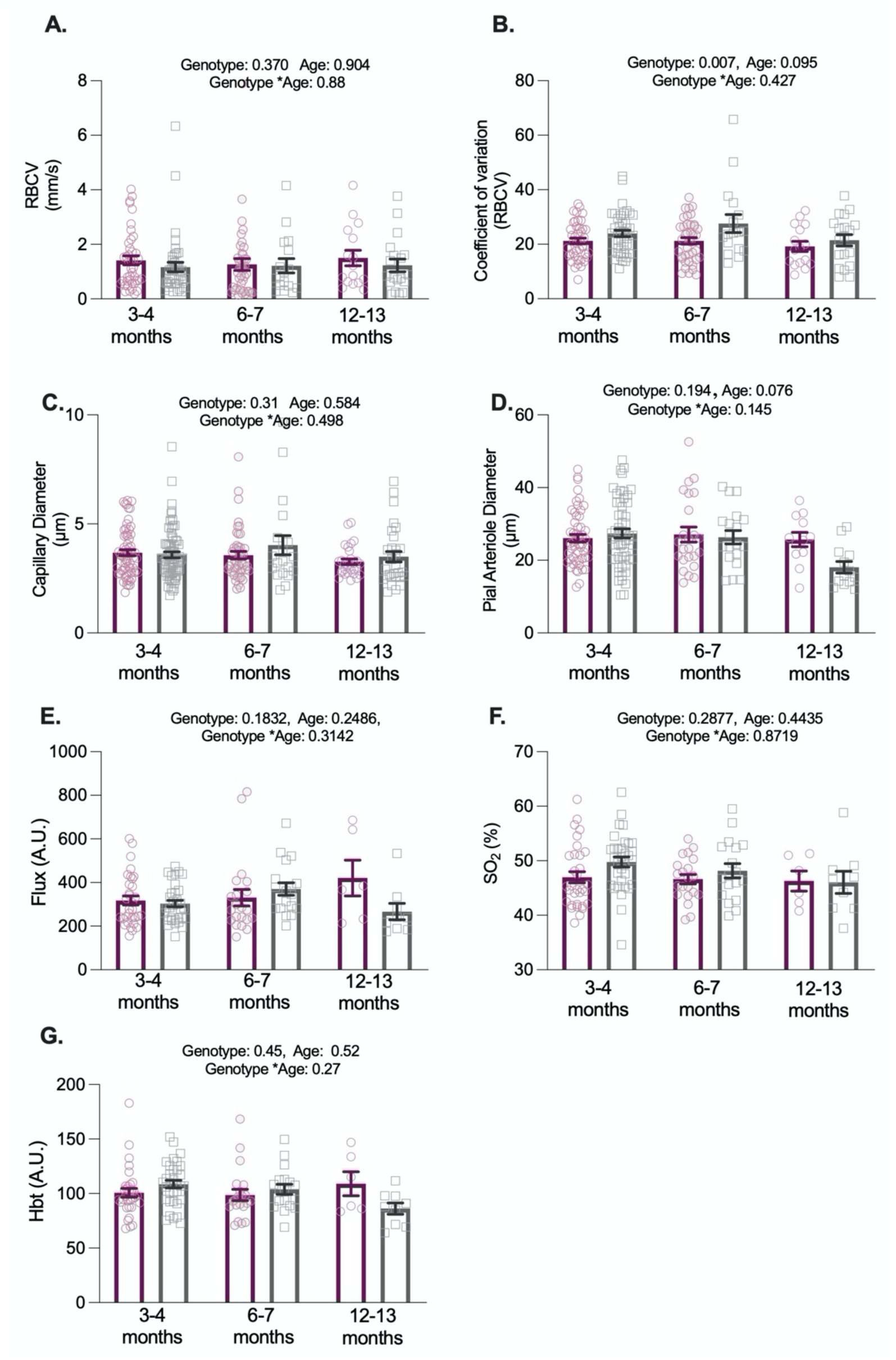
Under baseline conditions functional measurements are largely preserved in APOE4 mice within brain regions. This effect is preserved in older animals. There was no effect of genotype or age on **(A)** RBCV, **(C)** capillary diameter, **(D)** arteriole diameter, **(E)** flux, **(F)** SO_2_ or **(G)** HbT. However a significant effect of genotype was observed in RBCV CV measurements (B). Individual data points on bar graphs represent single vessels (A-D); or recording sessions (E-G). See appendix (i) for sample sizes and statistical tests.

## Notes

### Competing Interest Statement

The authors have declared no competing interest.

### Summary of Updates

This version of the paper includes data from the hippomcampus of APOE3 and APOE4 targeted replacement mice. All figures have therefore been revised to include these data. We have also simplified the analysis of vasomotion as well to aid readability.

## References

Aldea, R., Weller, R.O., Wilcock, D.M., Carare, R.O., Richardson, G., 2019. Cerebrovascular smooth muscle cells as the drivers of intramural periarterial drainage of the brain. Front. Aging Neurosci. 11, 1. doi:10.3389/fnagi.2019.00001

Angleys, H., Østergaard, L., Jespersen, S.N., 2015. The effects of capillary transit time heterogeneity (CTH) on brain oxygenation. J. Cereb. Blood Flow Metab. 35, 806–817. doi:10.1038/jcbfm.2014.254

Arganda-Carreras, I., Fernández-González, R., Muñoz-Barrutia, A., Ortiz-De-Solorzano, C., 2010. 3D reconstruction of histological sections: Application to mammary gland tissue. Microsc. Res. Tech. 73, 1019–1029. doi:10.1002/jemt.20829

Bell, R.D., Winkler, E.A., Singh, I., Sagare, A.P., Deane, R., Wu, Z., Holtzman, D.M., Betsholtz, C., Armulik, A., Sallstrom, J., Berk, B.C., Zlokovic, B.V., 2012. Apolipoprotein E controls cerebrovascular integrity via cyclophilin A. Nature 485, 512–516. doi:10.1038/nature11087

Blinder, P., Tsai, P.S., Kaufhold, J.P., Knutsen, P.M., Suhl, H., Kleinfeld, D., 2013. The cortical angiome: an interconnected vascular network with noncolumnar patterns of blood flow. Nat. Neurosci. 16, 889–897. doi:10.1038/nn.3426

Browndyke, J.N., Wright, M.C., Yang, R., Syed, A., Park, J., Hall, A., Martucci, K., Devinney, M.J., Shaw, L., Waligorska, T., Moretti, E.W., Whitson, H.E., Cohen, H.J., Mathew, J.P., Berger, M., MADCO-PC Investigators, 2021. Perioperative neurocognitive and functional neuroimaging trajectories in older APOE4 carriers compared with non-carriers: secondary analysis of a prospective cohort study. Br. J. Anaesth. 127, 917–928. doi:10.1016/j.bja.2021.08.012

Cai, Y., Hu, H., Liu, P., Feng, G., Dong, W., Yu, B., Zhu, Y., Song, J., Zhao, M., 2012. Association between the apolipoprotein E4 and postoperative cognitive dysfunction in elderly patients undergoing intravenous anesthesia and inhalation anesthesia. Anesthesiology 116, 84–93. doi:10.1097/ALN.0b013e31823da7a2

Chen, B.R., Kozberg, M.G., Bouchard, M.B., Shaik, M.A., Hillman, E.M.C., 2014. A critical role for the vascular endothelium in functional neurovascular coupling in the brain. J. Am. Heart Assoc. 3, e000787. doi:10.1161/JAHA.114.000787

Chen, T.-W., Wardill, T.J., Sun, Y., Pulver, S.R., Renninger, S.L., Baohan, A., Schreiter, E.R., Kerr, R.A., Orger, M.B., Jayaraman, V., Looger, L.L., Svoboda, K., Kim, D.S., 2013. Ultrasensitive fluorescent proteins for imaging neuronal activity. Nature 499, 295–300. doi:10.1038/nature12354

Cooper, C.E., 2003. Competitive, reversible, physiological? Inhibition of mitochondrial cytochrome oxidase by nitric oxide. IUBMB Life 55, 591–597. doi:10.1080/15216540310001628663

Corder, E.H., Saunders, A.M., Strittmatter, W.J., Schmechel, D.E., Gaskell, P.C., Small, G.W., Roses, A.D., Haines, J.L., Pericak-Vance, M.A., 1993. Gene dose of apolipoprotein E type 4 allele and the risk of Alzheimer’s disease in late onset families. Science 261, 921–923. doi:10.1126/science.8346443

Cumming, G., Finch, S., 2005. Inference by eye: confidence intervals and how to read pictures of data. Am. Psychol. 60, p170–180. doi:10.1037/0003-066X.60.2.170

Dana, H., Chen, T.-W., Hu, A., Shields, B.C., Guo, C., Looger, L.L., Kim, D.S., Svoboda, K., 2014. Thy1-GCaMP6 transgenic mice for neuronal population imaging in vivo. PLoS One 9, e108697. doi:10.1371/journal.pone.0108697

Das, A., Murphy, K., Drew, P.J., 2021. Rude mechanicals in brain haemodynamics: non-neural actors that influence blood flow. Philos. Trans. R. Soc. Lond. B, Biol. Sci. 376, 20190635. doi:10.1098/rstb.2019.0635

Deane, R., Sagare, A., Hamm, K., Parisi, M., Lane, S., Finn, M.B., Holtzman, D.M., Zlokovic, B.V., 2008. apoE isoform –specific disruption of amyloid β peptide clearance from mouse brain. J. Clin. Invest. 118, 4002–4013.

Di Marco, L.Y., Farkas, E., Martin, C., Venneri, A., Frangi, A.F., 2015. Is vasomotion in cerebral arteries impaired in alzheimer’s disease? J. Alzheimers Dis. 46, 35–53. doi:10.3233/JAD-142976

Drew, P.J., Blinder, P., Cauwenberghs, G., Shih, A.Y., Kleinfeld, D., 2010. Rapid determination of particle velocity from space-time images using the Radon transform. J. Comput. Neurosci. 29, 5–11. doi:10.1007/s10827-009-0159-1

Ganfield, R.A., Nair, P., Whalen, W.J., 1970. Mass transfer, storage, and utilization of O2 in cat cerebral cortex. Am. J. Physiol. 219, 814–821. doi:10.1152/ajplegacy.1970.219.3.814

Gould, I.G., Tsai, P., Kleinfeld, D., Linninger, A., 2017. The capillary bed offers the largest hemodynamic resistance to the cortical blood supply. J. Cereb. Blood Flow Metab. 37, 52–68. doi:10.1177/0271678X16671146

Greenberg, S.M., Bacskai, B.J., Hernandez-Guillamon, M., Pruzin, J., Sperling, R., van Veluw, S.J., 2020. Cerebral amyloid angiopathy and Alzheimer disease - one peptide, two pathways. Nat. Rev. Neurol. 16, 30–42. doi:10.1038/s41582-019-0281-2

Grijseels, D.M., Shaw, K., Barry, C., Hall, C.N., 2021. Choice of method of place cell classification determines the population of cells identified. BioRxiv. doi:10.1101/2021.02.26.433025

Hall, C.N., Reynell, C., Gesslein, B., Hamilton, N.B., Mishra, A., Sutherland, B.A., O’Farrell, F.M., Buchan, A.M., Lauritzen, M., Attwell, D., 2014. Capillary pericytes regulate cerebral blood flow in health and disease. Nature 508, 55–60. doi:10.1038/nature13165

Halliday, M.R., Rege, S.V., Ma, Q., Zhao, Z., Miller, C.A., Winkler, E.A., Zlokovic, B.V., 2016. Accelerated pericyte degeneration and blood-brain barrier breakdown in apolipoprotein E4 carriers with Alzheimer’s disease. J. Cereb. Blood Flow Metab. 36, 216–227. doi:10.1038/jcbfm.2015.44

Hamilton, N.B., Attwell, D., Hall, C.N., 2010. Pericyte-mediated regulation of capillary diameter: a component of neurovascular coupling in health and disease. Front. Neuroenergetics 2. doi:10.3389/fnene.2010.00005

Hawkes, C.A., Sullivan, P.M., Hands, S., Weller, R.O., Nicoll, J.A.R., Carare, R.O., 2012. Disruption of arterial perivascular drainage of amyloid-β from the brains of mice expressing the human APOE ε4 allele. PLoS One 7, e41636. doi:10.1371/journal.pone.0041636

He, Y., Wang, M., Chen, X., Pohmann, R., Polimeni, J.R., Scheffler, K., Rosen, B.R., Kleinfeld, D., Yu, X., 2018. Ultra-Slow Single-Vessel BOLD and CBV-Based fMRI Spatiotemporal Dynamics and Their Correlation with Neuronal Intracellular Calcium Signals. Neuron 97, 925–939.e5. doi:10.1016/j.neuron.2018.01.025

Henneman, W.J.P., Sluimer, J.D., Barnes, J., van der Flier, W.M., Sluimer, I.C., Fox, N.C., Scheltens, P., Vrenken, H., Barkhof, F., 2009. Hippocampal atrophy rates in Alzheimer disease: added value over whole brain volume measures. Neurology 72, 999–1007. doi:10.1212/01.wnl.0000344568.09360.31

Henson, R.N., Suri, S., Knights, E., Rowe, J.B., Kievit, R.A., Lyall, D.M., Chan, D., Eising, E., Fisher, S.E., 2020. Effect of apolipoprotein E polymorphism on cognition and brain in the Cambridge Centre for Ageing and Neuroscience cohort. Brain Neurosci. Adv. 4, 2398212820961704. doi:10.1177/2398212820961704

Iadecola, C., 2017. The Neurovascular Unit Coming of Age: A Journey through Neurovascular Coupling in Health and Disease. Neuron 96, 17–42. doi:10.1016/j.neuron.2017.07.030

Iadecola, C., Yang, G., Ebner, T.J., Chen, G., 1997. Local and propagated vascular responses evoked by focal synaptic activity in cerebellar cortex. J. Neurophysiol. 78, 651–659. doi:10.1152/jn.1997.78.2.651

Iturria-Medina, Y., Sotero, R.C., Toussaint, P.J., Mateos-Pérez, J.M., Evans, A.C., Alzheimer’s Disease Neuroimaging Initiative, 2016. Early role of vascular dysregulation on late-onset Alzheimer’s disease based on multifactorial data-driven analysis. Nat. Commun. 7, 11934. doi:10.1038/ncomms11934

Jiang, L., Zhong, J., Dou, X., Cheng, C., Huang, Z., Sun, X., 2015. Effects of ApoE on intracellular calcium levels and apoptosis of neurons after mechanical injury. Neuroscience 301, 375–383. doi:10.1016/j.neuroscience.2015.06.005

Jiang, Q., Lee, C.Y.D., Mandrekar, S., Wilkinson, B., Cramer, P., Zelcer, N., Mann, K., Lamb, B., Willson, T.M., Collins, J.L., Richardson, J.C., Smith, J.D., Comery, T.A., Riddell, D., Holtzman, D.M., Tontonoz, P., Landreth, G.E., 2008. ApoE promotes the proteolytic degradation of Abeta. Neuron 58, 681–693. doi:10.1016/j.neuron.2008.04.010

Juul Rasmussen, I., Rasmussen, K.L., Nordestgaard, B.G., Tybjærg-Hansen, A., Frikke-Schmidt, R., 2020. Impact of cardiovascular risk factors and genetics on 10-year absolute risk of dementia: risk charts for targeted prevention. Eur. Heart J. 41, p4024–4033. doi:10.1093/eurheartj/ehaa695

Kisler, K., Nelson, A.R., Montagne, A., Zlokovic, B.V., 2017. Cerebral blood flow regulation and neurovascular dysfunction in Alzheimer disease. Nat. Rev. Neurosci. 18, 419–434. doi:10.1038/nrn.2017.48

Kloske, C.M., Wilcock, D.M., 2020. The important interface between apolipoprotein E and neuroinflammation in alzheimer’s disease. Front. Immunol. 11, 754. doi:10.3389/fimmu.2020.00754

Knouff, C., Hinsdale, M.E., Mezdour, H., Altenburg, M.K., Watanabe, M., Quarfordt, S.H., Sullivan, P.M., Maeda, N., 1999. Apo E structure determines VLDL clearance and atherosclerosis risk in mice. J. Clin. Invest. 103, 1579–1586. doi:10.1172/JCI6172

Koizumi, K., Hattori, Y., Ahn, S.J., Buendia, I., Ciacciarelli, A., Uekawa, K., Wang, G., Hiller, A., Zhao, L., Voss, H.U., Paul, S.M., Schaffer, C., Park, L., Iadecola, C., 2018. Apoε4 disrupts neurovascular regulation and undermines white matter integrity and cognitive function. Nat. Commun. 9, 3816. doi:10.1038/s41467-018-06301-2

Kovacs-Oller, T., Ivanova, E., Bianchimano, P., Sagdullaev, B.T., 2020. The pericyte connectome: spatial precision of neurovascular coupling is driven by selective connectivity maps of pericytes and endothelial cells and is disrupted in diabetes. Cell Discov. 6, 39. doi:10.1038/s41421-020-0180-0

Lab, B.E., 2021. BrainEnergyLab/HCvsV1_NVC_Manuscript: NVCinHCmanuscript_March2021_release1. Zenodo. doi:10.5281/zenodo.4593010

Larramona-Arcas, R., González-Arias, C., Perea, G., Gutiérrez, A., Vitorica, J., García-Barrera, T., Gómez-Ariza, J.L., Pascua-Maestro, R., Ganfornina, M.D., Kara, E., Hudry, E., Martinez-Vicente, M., Vila, M., Galea, E., Masgrau, R., 2020. Sex-dependent calcium hyperactivity due to lysosomal-related dysfunction in astrocytes from APOE4 versus APOE3 gene targeted replacement mice. Mol. Neurodegener. 15, 35. doi:10.1186/s13024-020-00382-8

Lyons, D.G., Parpaleix, A., Roche, M., Charpak, S., 2016. Mapping oxygen concentration in the awake mouse brain. Elife 5. doi:10.7554/eLife.12024

Mateo, C., Knutsen, P.M., Tsai, P.S., Shih, A.Y., Kleinfeld, D., 2017. Entrainment of Arteriole Vasomotor Fluctuations by Neural Activity Is a Basis of Blood-Oxygenation-Level-Dependent “Resting-State” Connectivity. Neuron 96, 936–948.e3. doi:10.1016/j.neuron.2017.10.012

Mayhew, J.E., Askew, S., Zheng, Y., Porrill, J., Westby, G.W., Redgrave, P., Rector, D.M., Harper, R.M., 1996. Cerebral vasomotion: a 0.1-Hz oscillation in reflected light imaging of neural activity. Neuroimage 4, 183–193. doi:10.1006/nimg.1996.0069

Michaelis, E.K., 2012. Selective neuronal vulnerability in the hippocampus: relationship to neurological diseases and mechanisms for differential sensitivity of neurons to stress, in: Bartsch, T. (Ed.), The Clinical Neurobiology of the Hippocampus: An Integrative View. Oxford University Press, pp. 54–76. doi:10.1093/acprof:oso/9780199592388.003.0004

Miners, J.S., Van Helmond, Z., Chalmers, K., Wilcock, G., Love, S., Kehoe, P.G., 2006. Decreased expression and activity of neprilysin in Alzheimer disease are associated with cerebral amyloid angiopathy. J. Neuropathol. Exp. Neurol. 65, 1012–1021. doi:10.1097/01.jnen.0000240463.87886.9a

Montagne, A., Nation, D.A., Sagare, A.P., Barisano, G., Sweeney, M.D., Chakhoyan, A., Pachicano, M., Joe, E., Nelson, A.R., D’Orazio, L.M., Buennagel, D.P., Harrington, M.G., Benzinger, T.L.S., Fagan, A.M., Ringman, J.M., Schneider, L.S., Morris, J.C., Reiman, E.M., Caselli, R.J., Chui, H.C., Tcw, J., Chen, Y., Pa, J., Conti, P.S., Law, M., Toga, A.W., Zlokovic, B.V., 2020. APOE4 leads to blood-brain barrier dysfunction predicting cognitive decline. Nature 581, 71–76. doi:10.1038/s41586-020-2247-3

Najm, R., Jones, E.A., Huang, Y., 2019. Apolipoprotein E4, inhibitory network dysfunction, and Alzheimer’s disease. Mol. Neurodegener. 14, 24. doi:10.1186/s13024-019-0324-6

Nuriel, T., Angulo, S.L., Khan, U., Ashok, A., Chen, Q., Figueroa, H.Y., Emrani, S., Liu, L., Herman, M., Barrett, G., Savage, V., Buitrago, L., Cepeda-Prado, E., Fung, C., Goldberg, E., Gross, S.S., Hussaini, S.A., Moreno, H., Small, S.A., Duff, K.E., 2017. Neuronal hyperactivity due to loss of inhibitory tone in APOE4 mice lacking Alzheimer’s disease-like pathology. Nat. Commun. 8, 1464. doi:10.1038/s41467-017-01444-0

O’Donoghue, M.C., Murphy, S.E., Zamboni, G., Nobre, A.C., Mackay, C.E., 2018. APOE genotype and cognition in healthy individuals at risk of Alzheimer’s disease: A review. Cortex 104, 103–123. doi:10.1016/j.cortex.2018.03.025

Pachitariu, M., Stringer, C., Schröder, S., Dipoppa, M., Rossi, L.F., Carandini, M., Harris, K.D., 2016. Suite2p: beyond 10,000 neurons with standard two-photon microscopy. BioRxiv. doi:10.1101/061507

Parpaleix, A., Goulam Houssen, Y., Charpak, S., 2013. Imaging local neuronal activity by monitoring PO_2_ transients in capillaries. Nat. Med. 19, 241–246. doi:10.1038/nm.3059

Qiu, Z., Crutcher, K.A., Hyman, B.T., Rebeck, G.W., 2003. ApoE isoforms affect neuronal N-methyl-D-aspartate calcium responses and toxicity via receptor-mediated processes. Neuroscience 122, 291–303. doi:10.1016/j.neuroscience.2003.08.017

Royl, G., Füchtemeier, M., Leithner, C., Megow, D., Offenhauser, N., Steinbrink, J., Kohl-Bareis, M., Dirnagl, U., Lindasuer, U., 2008. Hypothermia effects on neurovascular coupling and cerebral metabolic rate of oxygen. Neuroimage 40, 1523–1532. doi:10.1016/j.neuroimage.2008.01.041

Schenning, K.J., Murchison, C.F., Mattek, N.C., Silbert, L.C., Kaye, J.A., Quinn, J.F., 2016. Surgery is associated with ventricular enlargement as well as cognitive and functional decline. Alzheimers Dement. 12, 590–597. doi:10.1016/j.jalz.2015.10.004

Shabir, O., Pendry, B., Lee, L., Eyre, B., Sharp, P., Rebollar, M.A., Howarth, C., Heath, P.R., Wharton, S.B., Francis, S.E., Berwick, J., 2020. Assessment of neurovascular coupling & cortical spreading depression in mixed models of atherosclerosis & alzheimer’s disease. BioRxiv. doi:10.1101/2020.08.13.249987

Shaw, K, Bell, L., Boyd, K., Grijseels, D.M., Clarke, D., Bonnar, O., Crombag, H.S., Hall, C.N., 2021a. Neurovascular coupling and oxygenation are decreased in hippocampus compared to neocortex because of microvascular differences. Nat. Commun. 12, 3190. doi:10.1038/s41467-021-23508-y

Shaw, Kira, Boyd, K., Anderle, S., Hammond-Haley, M., Amin, D., Bonnar, O., Hall, C.N., 2021b. Gradual not sudden change: multiple sites of functional transition across the microvascular bed. Front. Aging Neurosci. 13, 779823. doi:10.3389/fnagi.2021.779823

Sullivan, P.M., Mezdour, H., Aratani, Y., Knouff, C., Najib, J., Reddick, R.L., Quarfordt, S.H., Maeda, N., 1997. Targeted replacement of the mouse apolipoprotein E gene with the common human APOE3 allele enhances diet-induced hypercholesterolemia and atherosclerosis. J. Biol. Chem. 272, 17972–17980. doi:10.1074/jbc.272.29.17972

Sun, X., He, G., Qing, H., Zhou, W., Dobie, F., Cai, F., Staufenbiel, M., Huang, L.E., Song, W., 2006. Hypoxia facilitates Alzheimer’s disease pathogenesis by up-regulating BACE1 gene expression. Proc. Natl. Acad. Sci. USA 103, 18727–18732. doi:10.1073/pnas.0606298103

Thakore, P., Alvarado, M.G., Ali, S., Mughal, A., Pires, P.W., Yamasaki, E., Pritchard, H.A., Isakson, B.E., Tran, C.H.T., Earley, S., 2021. Brain endothelial cell TRPA1 channels initiate neurovascular coupling. Elife 10, e63040. doi:10.7554/eLife.63040

Thambisetty, M., Beason-Held, L., An, Y., Kraut, M.A., Resnick, S.M., 2010. APOE epsilon4 genotype and longitudinal changes in cerebral blood flow in normal aging. Arch. Neurol. 67, 93–98. doi:10.1001/archneurol.2009.913

Thomsen, M.S., Routhe, L.J., Moos, T., 2017. The vascular basement membrane in the healthy and pathological brain. J. Cereb. Blood Flow Metab. 37, 3300–3317. doi:10.1177/0271678X17722436

Uchida, K., Reilly, M.P., Asakura, T., 1998. Molecular stability and function of mouse hemoglobins. Zool Sci 15, 703–706. doi:10.2108/zsj.15.703

van Veluw, S.J., Hou, S.S., Calvo-Rodriguez, M., Arbel-Ornath, M., Snyder, A.C., Frosch, M.P., Greenberg, S.M., Bacskai, B.J., 2020. Vasomotion as a driving force for paravascular clearance in the awake mouse brain. Neuron 105, 549–561.e5. doi:10.1016/j.neuron.2019.10.033

Wierenga, C.E., Clark, L.R., Dev, S.I., Shin, D.D., Jurick, S.M., Rissman, R.A., Liu, T.T., Bondi, M.W., 2013. Interaction of age and APOE genotype on cerebral blood flow at rest. J. Alzheimers Dis. 34, 921–935. doi:10.3233/JAD-121897

Yamazaki, Y., Zhao, N., Caulfield, T.R., Liu, C.-C., Bu, G., 2019. Apolipoprotein E and Alzheimer disease: pathobiology and targeting strategies. Nat. Rev. Neurol. 15, 501–518. doi:10.1038/s41582-019-0228-7

Zhu, X., Bergles, D.E., Nishiyama, A., 2008. NG2 cells generate both oligodendrocytes and gray matter astrocytes. Development 135, 145–157. doi:10.1242/dev.004895

Zlokovic, B.V., 2011. Neurovascular pathways to neurodegeneration in Alzheimer’s disease and other disorders. Nat. Rev. Neurosci. 12, 723–738. doi:10.1038/nrn3114

Zott, B., Simon, M.M., Hong, W., Unger, F., Chen-Engerer, H.-J., Frosch, M.P., Sakmann, B., Walsh, D.M., Konnerth, A., 2019. A vicious cycle of β amyloid-dependent neuronal hyperactivation. Science 365, 559–565. doi:10.1126/science.aay0198

