## Appendix containing full statistics reports for all analyses for "APOE4 expression confers a mild, persistent reduction in neurovascular function in the visual cortex and hippocampus of awake mice"

| **Figure label** | **n** | **Mean**  Appendix (i)  Figure 1 | **Standard Deviation** | **Test** | **Test Statistic**  **F value**  G = Genotype  BR = Brain Region | **P Value** |
| --- | --- | --- | --- | --- | --- | --- |
| 1B | V1 E3: 16/3  V1 E4: 19/3  HC E3: 23/3  HC E4: 23/3  Slices/Animals | V1 E3: 2.638  V1 E4: 2.133  HC E3: 1.901  HC E4: 0.928 | V1 E3: 0.784  V1 E4: 0.463  HC E3: 0.733  HC E4: 0.344 | Linear mixed model (LMER function (R studio))  Random factor = Animal ID | G: 9.379  BR: 62.9534  G * BR: 1.768 | G: 0.04374  BR: 1.611e-11  G * BR: 0.18771 |
| 1E | V1 E3: 42/12  V1 E4: 41/11  HC E3: 37/7  HC E4: 26/6  Vessels/Animals | V1 E3: 1.415  V1 E4: 1.168  HC E3: 1.461  HC E4: 1.745 | V1 E3: 1.05  V1 E4: 1.128  HC E3: 0.553  HC E4: 0.936 | Linear mixed model  (LMER function (R studio))  Random factor = Animal ID | G: 0.0038  BR: 2.2247  G * BR: 1.9206 | G: 0.9513  BR: 0.1525  G * BR: 0.1821 |
| 1F | V1 E3: 42/12  V1 E4: 41/11  HC E3: 37/7  HC E4: 26/6  Vessels/Animals | V1 E3: 21.19  V1 E4: 23.93  HC E3: 16.191  HC E4: 20.944 | V1 E3: 6.829  V1 E4: 7.741  HC E3: 7.616  HC E4: 10.92 | Linear mixed model  (LMER function (R studio))  Random factor = Animal ID | G: 6.327  BR: 7.481  G * BR: 0.516 | G: 0.002  BR: 0.012  G * BR: 0.480 |
| 1G | V1 E3: 29/11  V1 E4: 30/11  HC E3: 37/7  HC E4: 26/6  Vessels/Animals | V1 E3: 54.253  V1 E4: 49.769  HC E3: 57.880  HC E4: 52.961 | V1 E3: 14.204  V1 E4: 13.508  HC E3: 7.046  HC E4: 6.555 | Linear mixed model  (LMER function (R studio))  Random factor = Animal ID | G: 3.6886  BR: 1.6496  G * BR: 0.0107 | G: 0.06752  BR:0.21205  G * BR:0.91865 |
| 1H | V1 E3: 64/13  V1 E4:72/13  HC E3: 37/7  HC E4: 26/6  Vessels/Animals | V1 E3: 3.679  V1 E4: 3.582  HC E3: 3.543  HC E4: 3.8247 | V1 E3: 1.1188841  V1 E4: 1.1724786  HC E3: 0.9076  HC E4: 0.78756 | Linear mixed model  (LMER function (R studio))  Random factor = Animal ID | G:0.2811  BR: 0.0005  G * BR:0.1185 | G:0.5997  BR:0.9817  G * BR:0.7330 |
| 1I | V1 E3: 32/15  V1 E4: 33/15  HC E3: 19/8  HC E4: 18/8  Sessions/Animals | V1 E3: 317.5321  V1 E4: 303.4218  HC E3: 221.1394  HC E4: 184.1300 | V1 E3: 117.59148  V1 E4: 86.64690  HC E3: 99.22186  HC E4: 112.96310 | Linear mixed model  (LMER function (R studio))  Random factor = Animal ID | G: 1.2208  BR: 17.9543  G * BR: 0.0639 | G:0.2758363  BR:0.0001303  G * BR: 0.8017816 |
| 1J | V1 E3: 32/15  V1 E4: 33/15  HC E3: 19/8  HC E4: 18/8  Sessions/Animals | V1 E3: 100.78365  V1 E4: 108.66546  HC E3: 32.23938  HC E4: 41.28505 | V1 E3: 22.739420  V1 E4: 20.320726  HC E3: 9.807566  HC E4: 14.169297 | Linear mixed model  (LMER function (R studio))  Random factor = Animal ID | G :2.9684  BR: 224.4322  G * BR:0.0827 | G: 0.09415  BR: 2e-16  G * BR:0.77549 |
| 1K | V1 E3: 32/15  V1 E4: 33/15  HC E3: 19/8  HC E4: 18/8  Sessions/Animals | V1 E3: 46.97072  V1 E4: 49.78518  HC E3: 23.52394  HC E4: 26.15068 | V1 E3: 5.777214  V1 E4: 5.344955  HC E3: 7.332528  HC E4: 6.757558 | Linear mixed model  (LMER function (R studio))  Random factor = Animal ID | G:3.3477  BR: 247.3893  G * BR: 0.0429 | G: 0.07604  BR:2e-16  G * BR: 0.83707 |
| 1L | V1 E3: 32/15  V1 E4: 33/15  HC E3: 19/8  HC E4: 18/8  Sessions/Animals | V1 E3: 167.3166  V1 E4: 151.6007  HC E3: 166.9108  HC E4: 132.7893 | V1 E3: 63.80262  V1 E4: 44.95330  HC E3: 73.81183  HC E4: 75.29335 | Linear mixed model  (LMER function (R studio))  Random factor = Animal ID | G:2.4841  BR:1.0570  G * BR: 0.2096 | G:0.1226  BR:0.3099  G * BR:0.6495 |

| **Figure label** | **n**  Figure 2 | **Mean** | **Standard Deviation** | **Test** | **Test Statistic**  **F value**  G = Genotype  BR = Brain Region | **P Value** |
| --- | --- | --- | --- | --- | --- | --- |
| 2C | V1 E3: 35/6  V1 E4: 33/5  HC E3: 41/6  HC E4: 40/7  FOVs/Animals | V1 E3: 0.0438  V1 E4: 0.0381  HC E3: 0.0251  HC E4: 0.0715 | V1 E3:0.0419  V1 E4:0.0307    HC E3:0.04323244  HC E4: 0.1213 | Linear mixed model (LMER function (R studio))  Random factor = Animal ID | G:0.5088  BR: 0.0706  G * BR: 1.0297 | G: 0.4842  BR: 0.7932  G * BR: 0.3227 |
| 2D | V1 E3: 1715/6  V1 E4: 1635/5  HC E3: 2571/6  HC E4: 2705/7  Cells/Animals | V1 E3: 2.2597  V1 E4: 2.083  HC E3: 2.5491  HC E4: 2.6826 | V1 E3:3.4178  V1 E4:3.0133    HC E3:2.8967  HC E4:3.3684 | Linear mixed model  (LMER function (R studio))  Random factor = Animal ID | G:0.0238  BR: 1.7821  G * BR: 0.1742 | G: 0.8791  BR: 0.1973  G * BR: 0.6810 |
| 2E | V1 E3: 1768/6  V1 E4: 1676/5  HC E3: 3411/6  HC E4: 2970/7  Cells/Animals | V1 E3: 5.5935  V1 E4: 5.5769    HC E3: 3.6409  HC E4: 3.5793 | V1 E3: 3.6658  V1 E4:3.4444    HC E3:4.1667  HC E4:3.2322 | Linear mixed model  (LMER function (R studio))  Random factor = Animal ID | G:0.3695  BR: 5.7217  G * BR: 0.0176 | G: 0.5536  BR: 0.0323  G * BR: 0.8963 |
| 2G | V1 E3: 6/503  V1 E4: 6/593  Animals/Cells | V1 E3: 0.9394  V1 E4: 1.428 | V1 E3: 0.1979  V1 E4: 0.4092 | Mann-Whitney U | MWU = 2 | P = 0.0087 |
| 2H | V1 E3: 6  V1 E4: 6  Animals | V1 E3: 24.89  V1 E4: 25.16 | V1 E3: 4.472  V1 E4: 7.922 | T-test | T = 0.07084 | P = 0.9449 |

Figure 3

| **Figure label** | **n** | **Mean** | **Standard Deviation** | **Test** | **Test Statistic** | **P Value** |
| --- | --- | --- | --- | --- | --- | --- |
| 3C | E3: 59/15  E4: 61/15  Vessels/Animals | E3: 21.97  E4: 23.19 | V1 E3: 14.11  V1 E4: 14.43 | MWU | U = 1685 | 0.55 |
| 3D | E3: 60/15  E4: 64/15  Vessels/Animals | E3: 70.2  E4: 61.97 | V1 E3:26.05  V1 E4: 26.66 | Weighted Least Squares linear regression.  Weight: number of trials | Power = -0.5 | 0.034 |
| 3G | E3: 51/12  E4: 62/13  Vessels/Animals | E3: 17.88  E4: 19.68 | V1 E3:21.05  V1 E4: 15.71 | MWU | U = 1350 | 0.1843 |
| 3H | E3: 60/13  E4: 71/13  Vessels/Animals | E3:37.87  E4: 38.09 | V1 E3:27.73  V1 E4: 28.92 | Weighted Least Squares linear regression.  Weight: number of trials | Power = -0.5 | 0.328 |
| 3K | E3: 33/12  E4:27/11  Vessels/Animals | E3:96.38  E4:99.94 | V1 E3:66.48  V1 E4: 100.7 | MWU | U = 423 | 0.7465 |
| 3L | E3: 39/12  E4: 37/11  Vessels/Animals | E3: 44.96  E4: 34.83 | V1 E3: 30.5  V1 E4: 30.59 | Weighted Least Squares linear regression.  Weight: number of trials | Power = -1 | 0.018 |

| **Figure label**  Figure 4 | **n** | **Mean** | **Standard Deviation** | **Test** | **Test Statistic** | **P Value** |
| --- | --- | --- | --- | --- | --- | --- |
| 4A | HC E3: 43/6  HC E4: 72/7  Vessels/Animals | HC E3:26.24  HC E4:26.33 | HC E3:21.08  HC E4: 28.48 | Weighted Least Squares linear regression.  Weight: number of trials | Power = -0.05 | P = 0.89 |
| 4B | HC E3: 43/6  HC E4: 72/7  Vessels/Animals | HC E3: 16.28 /83.72  HC E4: 34.72/ 65.278    Non-Responsive/  Responsive | N/A | Chi squared | Chi squared = 4.56 | P = 0.03 |
| 4C | HC E3: 422/6  HC E4: 584/7  Events/Animals | HC E3:24.17/  75.829  HC E4:14.384/ 85.617    Non-Responsive/  Responsive | N/A | Chi squared | Chi squared = 15.57 | P < 0.0001 |
| 4E | HC E3: 102/6  HC E4: 84/7  Events/Animals | HC E3: 0.1924  HC E4:0.2294 | HC E3:0.172  HC E4: 0.2041 | Linear mixed model  (LMER function (R studio))  Random factor = Animal ID | 0.8695 | P = 0.3827 |
| 4G | HC E3: 102/6  HC E4: 84/7  Events/Animals | HC E3: 0.0337  HC E4:0.04762 | HC E3:0.0648  HC E4: 0.1119 | Linear mixed model(LMER function (R studio))  Random factor = Animal ID | 0.6245 | 0.4502 |
| 4H NVC | HC E3: 416/6  HC E4: 527/7  Events/Animals | HC E3: 0.053  HC E4: -0.844 | HC E3:1.306  HC E4: 19.71 | Linear mixed model  (LMER function (R studio))  Random factor = Animal ID | 0.8602 | 0.3539 |

Figure 5

| **Figure label** | **n** | **Mean** | **Standard Deviation** | **Test** | **Test Statistic** | **P Value** |
| --- | --- | --- | --- | --- | --- | --- |
| 5E | E3: 52/15  E4: 53/15  Vessels/animals | E3: 5.967  E4: 3.754 | E3: 5.651  E4:3.449 | Mann-Whitney U | U = 1054 | P = 0.0373 |
| 5F | E3: 52/15  E4: 53/15  Vessels/animals | N/A |  | Spearman r | E3 r = 0.1589  E4 r = -0.052 | E3 = 0.2606  E4 = 0.714 |
| 5H | E3: 1770/6  E4: 1677/5  Cellss/Animals | E3: 4.251  E4: 22.48 | E3: 3.925  E4: 29.71 | Welch’s T test | T = 1.362 | P = 0.2429 |

Figure 6

| **Figure label** | **n**  Y = 3-4mo  M = 6-7 mo  O – 12-13mo | **Mean** | **Standard Deviation** | **Test** | **Test Statistic**  **F value**  G = Genotype  A = Age | **P Value** |
| --- | --- | --- | --- | --- | --- | --- |
| 6D | E3 Y: 59/15  E4 Y: 61/15  E3 M: 35/10  E4 M:31/10  E3 O: 13/4  E4 O: 16/4  Vessels/animals | E3 Y: 21.97  E4 Y: 23.19  E3 M: 13.02  E4 M: 19.87  E3 O: 19.27  E4 O: 16.54 | E3 Y: 14.11  E4 Y: 14.43  E3 M: 7.136  E4 M: 15.15  E3 O: 10.52  E4 O: 37.36 | Linear mixed model (LMER function (R studio))  Random factor = Animal ID | G: 0.0892  A: 4.411  G*A: 1.097 | G: 0.7683  A: 0.0183  G*A: 0.3432  Y-M: 0.0159  Y-O: 0.3378  M-O: 0.9993 |
| 6E | E3 Y: 33/12  E4 Y:27/11  E3 M: 23/8  E4 M:7/6  E3 O: 14/4  E4 O: 14/4  Vessels/animals | E3 Y: 96.38  E4 Y: 99.94  E3 M: 64.6  E4 M: 93.61  E3 O: 100.3  E4 O: 110.3 | E3 Y: 66.48  E4 Y: 100.7  E3 M: 47.75  E4 M: 48.54  E3 O: 53.01  E4 O: 57.45 | Linear mixed model (LMER function (R studio))  Random factor = Animal ID | G: 2.5737  A: 4.0378  G*A: 0.995 | G: 0.11576  A: 0.02137  G*A: 0.37428  Y-M: 0.0170  Y-O: 0.5434  M-O: 0.3779 |
| 6F | E3 Y: 60/13  E4 Y: 71/13  E3 M: 47/9  E4 M:15/7  E3 O: 29/4  E4O: 28/4  Vessels/animals | E3 Y: 37.87  E4 Y: 38.09  E3 M: 37.54  E4 M: 28.07  E3 O: 36.23  E4 O: 46.7 | E3 Y: 27.73  E4 Y: 28.92  E3 M: 19.14  E4 M: 28.71  E3 O:24.42  E4 O: 28.18 | Linear mixed model (LMER function (R studio))  Random factor = Animal ID | G: 0.8149  A: 0.7816  G*A:0.2325 | G: 0.3740  A: 0.4641  G*A: 0.7936 |
| 6G | E3 Y: 51/12  E4 Y: 62/13  E3 M: 44/9  E4 M:10/7  E3 O: 27/4  E4 O: 27/4  Vessels/animals | E3 Y: 17.88  E4 Y:19.68  E3 M: 9.971  E4 M: 11.31  E3 O: 10.29  E4 O: 20.54 | E3 Y: 21.05  E4 Y: 15.71  E3 M: 7.60  E4 M: 7.814  E3 O: 6.465  E4O: 17.14 | Weighted Least Squares linear regression.  Weight: number of trials | Power value: -0.5 | G: 0.033  A: 0.465  Regression: 0.08 |
| 6H | E3 Y: 60/15  E4 Y: 64/15  E3 M: 37/10  E4 M:33/10  E3 O: 13/4  E4 O: 16/4  Vessels/animals | E3 Y: 70.2  E4 Y: 61.97  E3 M: 60.38  E4 M: 54.97  E3 O: 78.7  E4 O: 71.2 | E3 Y: 26.05  E4 Y: 26.66  E3 M: 22.54  E4 M: 33.22  E3 O: 19.00  E4 O: 30.10 | Weighted Least Squares linear regression.  Weight: number of trials | Power value: -0.5 | G: 0.683  A:0.590  Regression: 0.777 |
| 6I | E3 Y: 39/12  E4 Y: 37/11  E3 M: 34/9  E4 M: 12/6  E3 O: 16/4  E4O: 17/4  Vessels/animals | E3 Y: 44.96  E4 Y: 34.83  E3 M: 35.33  E4 M: 22.83  E3 O: 44.97  E4O: 53.4 | E3 Y: 30.50  E4 Y: 30.59  E3 M: 35.66  E4 M: 29.01  E3 O: 28.71  E4O: 37.58 | Weighted Least Squares linear regression.  Weight: number of trials | Power value: -1 | G: 0.15  A: 0.648  Regression: 0.313 |
| 6K | E3 Y: 503/6  E4 Y:593 /6  E3 M: 171/5  E4 M: 136/3  Cells/animals | E3 Y: 0.9394  E4 Y: 1.428  E3 M: 0.7972  E4M: 1.625 | E3 Y: 0.1979  E4 Y: 0.44092  E3 M: 0.1970  E4 M: 0.9413 | Linear mixed model (LMER function (R studio))  Random factor = Animal ID | G: 7.9478  A: 0.0008  G*A: 1.3313 | G: 0.01679  A: 0.97794  G*A: 0.28226 |
| 6L | E3 Y: 1943/6  E4 Y: 2140/6  E3 M:666/5  E4 M:508/3  Cells/animals | E3 Y: 24.89  E4 Y: 25.16  E3 M: 26.8  E4 M: 25.86 | E3 Y: 4.472  E4 Y: 7.922  E3 M: 17.54  E4 M: 6.900 | Linear mixed model (LMER function (R studio))  Random factor = Animal ID | G: 0.0048  A: 0.0719  G*A: 0.0153 | G: 0.9455  A: 0.7920  G*A: 0.9031 |

| **Figure label** | **n**  Y = 3-4mo  M = 6-7 mo  O – 12-13mo | **Mean** | **Standard Deviation** | **Test** | **Test Statistic**  **F value**  G = Genotype  A = Age | **P Value** |
| --- | --- | --- | --- | --- | --- | --- |
| 7D | E3 Y: 52/15  E4 Y: 53/15  E3 M: 25/10  E4M: 18/8  E3 O: 12/4  E4O: 12/3  Vessels/animals | E3 Y: 5.967  E4 Y:3.754  E3 M: 5.510  E4M: 3.219  E3 O:5.239  E4O: 1.198 | E3 Y: 5.651  E4 Y: 3.449  E3 M: 4.251  E4M:3.414  E3 O:4.316  E4O:0.8474 | Linear mixed model (LMER function (R studio))  Random factor = Animal ID | G: 12.631  A: 1.366  G*A: 0.6723 | G: 0.00097  A: 0.26  G*A: 0.6723 |

```
          Figure 7


Figure 3:  Supplementary Figure 1
```

| **Figure label** | **n**  Y = 3-4mo  M = 6-7 mo | **Mean** | **Standard Deviation** | **Test** | **Test Statistic**  **F value**  G = Genotype  A = Age | **P Value** |
| --- | --- | --- | --- | --- | --- | --- |
| S2A | E3Y: 60/15  E4Y: 64/15  E3M: 37/10  E4M: 33/10  Vessels/Animals | E3Y: 13.99  E4Y: 8.785  E3M: 13.99  E4M: 8.785 | E3Y: 16.84  E4Y: 10.557  E3M: 9.289  E4M: 8.772 | 2 x 2 ANOVA | G: 5.508  A: 8.561  G*A: 0.201 | G: 0.02  A:0.004  G*A: 0.654 |
| S2B | E3Y: 60/13  E4Y: 71/13  E3M: 47/9  E4M:15/7  Vessels/Animals | E3: 6.428  E4: 2.019  E3M: 1.069  E4M: -0.773 | E3Y: 18.46  E4Y: 16.58  E3M: 10.60  E4M: 6.707 | 2 x 2 ANOVA | G: 1.378  A: 2.344  G*A: 0.232 | G: 0.242  A: 0.127  G*A: 0.630 |
| S2C | E3Y: 39/12  E4Y:37/11  E3M:34/9  E4M:12/6  Vessels/Animals | E3Y: 51.51  E4Y: 24.69  E3M: 22.04  E4M: 12.40 | E3Y: 64.94  E4Y: 52.60  E3M: 64.77  E4M: 24.68 | 2 x 2 ANOVA | G: 2.341  A:3.071  G*A: 0.520 | G: 0.129  A: 0.082  G*A:0.472 |

Figure 3: Supplementary Figure 2

| **Figure label** | **n** | **Mean** | **Standard Deviation** | **Test** | **Test Statistic** | **P Value** |
| --- | --- | --- | --- | --- | --- | --- |
| S3A | E3: 15  E4:15  Animals | E3: 19.38  E4: 19.17 | E3: 12.61  E4: 16.52 | T-Test | T = 0.03848 | P = 0.9696 |
| S3B | E3: 15  E4:15  Animals | E3: 10.8  E4: 9.1 | E3: 7.860  E4:10.07 | T-test | T = 0.5154 | P = 0.6103 |
| S3C | E3: 15  E4:15  Animals | E3: 8.026  E4: 7.291 | E3: 5.091  E4:5.887 | T-test | T = 0.3655 | P = 0.7175 |
| S3D | E3: 15  E4:15  Animals | E3: 7.561  E4: 7.246 | E3: 8.510  E4:6.793 | T-test | t= 0.1119 | P = 0.9117 |

| **Figure label** | **n** | **Mean** | **Standard Deviation** | **Test** | **Test Statistic** | **P Value** |
| --- | --- | --- | --- | --- | --- | --- |
| S4A | E3:538/8  E4:549/8  Events/Animals | E3:14.53  E4:21.00 | E3:15.92  E4:33.51 | Linear mixed model  (LMER function (R studio))  Random factor = Animal ID | 2.1297 | 0.1678 |
| S4B | E3:538/8  E4:549/8  Events/Animals | E3:37.10  E4:41.68 | E3:34.10  E4:46.84 | Linear mixed model  (LMER function (R studio))  Random factor = Animal ID | 1.5147 | 0.2406 |
| S4C | E3:538/8  E4:549/8  Events/Animals | E3: 7.835  E4: 7.281 | E3: 8.816  E4: 9.233 | Linear mixed model  (LMER function (R studio))  Random factor = Animal ID | 0.6893 | 0.4205 |
| S4D | E3:538/8  E4:549/8  Events/Animals | E3: 1.031  E4: 0.9946 | E3:1.127  E4:1.697 | Linear mixed model  (LMER function (R studio))  Random factor = Animal ID | 0.2449 | 0.6282 |
| S4E | E3:538/8  E4:549/8  Events/Animals | E3: 0.1433  E4: 0.1216 | E3: 0.2286  E4: 0.2327 | Linear mixed model  (LMER function (R studio))  Random factor = Animal ID | 1.6368 | 0.2234 |

Figure 4: Supplementary Figure 1

| **Figure label** | **n** | **Mean** | **Standard Deviation** | **Test** | **Test Statistic** | **P Value** |
| --- | --- | --- | --- | --- | --- | --- |
| S5A | E3:55/6  E4:57/7 | E3:8.096  E4:8.664 | E3:4.509  E4:5.605 | MWU | U= 1491 | 0.6595 |
| S5C | E3:54/6  E4:55/7 | E3:0.1832  E4:0.1729 | E3:0.3178  E4:0.2421 | MWU | U = 1350 | 0.4165 |
| S5F | E3:63/13  E4:71/13 | E3:0.05814  E4:0.07079 | E3:0.052  E4:0.078 | MWU | U = 2051 | 0.4108 |

Figure 5: Supplementary Figure 1

Figure 6: Supplementary Figure 1

| **Figure label** | **n** | **Mean** | **Standard Deviation** | **Test** | **Test Statistic**  **F value**  G = Genotype  A = Age | **P Value** |
| --- | --- | --- | --- | --- | --- | --- |
| S6A | E3 Y: 42/12  E4 Y: 41/11  E3 M: 40/9  E4M:17/6  E3 O: 16/4  E4O: 19/4  Vessels/animals | E3 Y: 1.415  E4 Y: 1.168  E3 M: 1.263  E4M: 1.215  E3 O: 1.501  E4O: 1.23 | E3Y: 1.05  E4Y: 1.128  E3M: 1.355  E4M: 1.069  E3O: 1.141  E4O: 1.007 | Linear mixed model (LMER function (R studio))  Random factor = Animal ID | G: 0.8406  A: 0.1009  G*A: 0.1275 | G: 0.370  A: 0.904  G*A: 0.8806 |
| S6B | E3 Y: 42/12  E4 Y: 41/11  E3 M: 40/9  E4M:17/6  E3 O: 16/4  E4O: 19/4  Vessels/animals | E3 Y: 21.19  E4 Y: 23.93  E3 M: 21.23  E4M: 27.57  E3 O: 19.19  E4O: 21.46 | E3Y: 6.829  E4Y: 7.741  E3M: 7.703  E4M: 13.55  E3O:7.44  E4O: 9.001 | Linear mixed model (LMER function (R studio))  Random factor = Animal ID | G: 7.4186  A: 2.3827  G*A: 0.8558 | G: 0.007  A: 0.0954  G*A: 0.427 |
| S6C | E3 Y: 64/13  E4 Y: 72/13  E3 M: 47/9  E4M:20/7  E3 O: 29/4  E4O: 30/4  Vessels/animals | E3 Y: 3.679  E4 Y: 3.582  E3 M: 3.568  E4M: 4.027  E3 O: 3.269  E4O: 3.497 | E3Y: 1.119  E4Y: 1.172  E3M: 1.177  E4M: 1.982  E3O: 0.6878  E4O: 1.318 | Linear mixed model (LMER function (R studio))  Random factor = Animal ID | G: 1.0766  A: 0.5427  G*A: 0.7056 | G: 0.308  Age: 0.5841  G*A: 0.498 |
| S6D | E3 Y: 52/15  E4 Y: 55/15  E3 M: 24/10  E4M: 19/8  E3 O: 12/4  E4O: 13/4  Vessels/animals | E3 Y: 26.06  E4 Y: 27.39  E3 M: 27.08  E4M: 26.32  E3 O: 25.73  E4O: 18.07 | E3Y: 7.589  E4Y: 9.337  E3M: 10.21  E4M: 8.007  E3O: 6.792  E4O: 5.706 | Linear mixed model (LMER function (R studio))  Random factor = Animal ID | G:1.744  A: 2.6679  G*A: 1.9831 | G: 0.194  A: 0.076  G*A: 0.145 |
| S6E | E3 Y: 32/15  E4 Y: 33/15  E3 M: 21/10  E4M:18/8  E3 O: 6/3  E4O: 9/4  Sessions/animal | E3 Y: 317.5  E4 Y: 303.4  E3 M: 330.6  E4M: 370.4  E3 O: 421  E4O: 266.6 | E3Y: 117.6  E4Y: 86.65  E3M: 174.2  E4M: 122.4  E3O: 201.1  E4O: 112.4 | Linear mixed model (LMER function (R studio))  Random factor = Animal ID | G:1.8496  A: 1.4343  G*A: 1.1799 | G: 0.1832  A: 0.2486  G*A: 0.3142 |
| S6F | E3 Y: 32/15  E4 Y: 33/15  E3 M: 21/10  E4M:18/8  E3 O: 6/3  E4O: 9/4  sessions/animal | E3 Y: 46.97  E4 Y: 49.79  E3 M: 46.62  E4M: 48.15  E3 O: 46.28  E4O: 46.03 | E3Y: 5.777  E4Y: 5.345  E3M: 4.010  E4M: 5.526  E3O:4.506  E4O:6.116 | Linear mixed model (LMER function (R studio))  Random factor = Animal ID | G: 1.1683  A: 0.8245  G*A: 0.1374 | G: 0.2877  A: 0.4435  G*A: 0.8719 |
| S6G | E3 Y: 32/15  E4 Y: 33/15  E3 M: 21/10  E4M:18/8  E3 O: 6/3  E4O: 9/4  sessions/animal | E3 Y: 100.8  E4 Y: 108.7  E3 M: 98.67  E4M: 103.9  E3 O: 109  E4O: 86.31 | E3Y: 22.74  E4Y: 20.32  E3M: 23.57  E4M: 19.54  E3O: 27.08  E4O: 15.55 | Linear mixed model (LMER function (R studio))  Random factor = Animal ID | G: 0.5867  A: 0.6544  G*A: 1.3212 | G: 0.448  A: 0.5233  G*A: 0.2743 |
